# A macrophage-derived noncanonical WNT niche drives immune exclusion in pancreatic cancer

**DOI:** 10.64898/2026.02.05.704114

**Authors:** Na Hyun Kim, Young Min Song, San Sung Kwon, Sang Hyub Lee, Eun Na Kim, Jung Joo Hong, Seung Hyeok Seok, Yi Rang Na

## Abstract

**Background and Aims:** Pancreatic ductal adenocarcinoma (PDAC) is characterized by profound immune exclusion and resistance to immunotherapy. Although WNT signaling has been implicated in PDAC, its cellular source within the tumor microenvironment and its contribution to immune suppression remain poorly defined. This study investigated whether myeloid-derived WNT signaling promotes PDAC progression.

**Methods:** Transcriptomic data from human PDAC cohorts, including The Cancer Genome Atlas (TCGA), and published single-cell RNA sequencing datasets were analyzed. Macrophage-associated WNT5A expression in human PDAC biopsies was assessed using in situ hybridization and immunofluorescence. Macrophage-derived WNT secretion was genetically disrupted using macrophage-specific *Porcn* knockout mice in orthotopic and subcutaneous KPC tumor models. Lineage-resolved spatial organization of macrophage subsets was characterized using Ms4a3 fate-mapping double-reporter mice with immunofluorescence and imaging mass cytometry. Macrophages–CD8⁺ T cells interactions were assessed using tumor-educated macrophage conditioned media, pharmacologic ARG1 inhibition, and in vivo CD8⁺ T cell depletion.

**Results:** PDAC tumors with high macrophage signatures showed enrichment of noncanonical WNT signaling, and macrophage-associated WNT5A was detected in human biopsies. Disruption of macrophage-derived WNT secretion suppressed tumor growth, reversed immune exclusion, and enhanced cytotoxic CD8⁺ T cell infiltration. Spatial lineage-resolved analysis demonstrated progressive accumulation of Hexb⁺ tissue-resident macrophages that dominated advanced lesions and formed a WNT-rich niche closely associated with Trem2⁺Arg1⁺ monocyte-derived macrophages. Mechanistically, macrophage-derived noncanonical WNT activated a JNK/c-Jun-ARG1 axis that inhibited CD8⁺ T cell proliferation, an effect abolished by myeloid WNT loss.

**Conclusions:** Myeloid-derived noncanonical WNT establishes a lineage-structured macrophage niche that enforces immune exclusion in PDAC. Targeting macrophage-restricted WNT signaling represents a promising strategy to reprogram the PDAC immune microenvironment.

## Introduction

Pancreatic ductal adenocarcinoma (PDAC) is among the most aggressive and treatment-refractory malignancies, with a five-year survival rate of approximately 11% ^1^. With a steadily increasing incidence, PDAC is project to become the second leading cause of cancer-related deaths in the United States by 2030 ^2^. Despite sustained therapeutic efforts, meaningful clinical progress in PDAC remains limited. Only 15–20% of patients are eligible for upfront curative resection, as most present with unresectable locally advanced or metastatic disease. Consequently, systemic chemotherapy remains the cornerstone of treatment across neoadjuvant, adjuvant, and palliative settings, most commonly using FOLFIRINOX or gemcitabine plus nab-paclitaxel ^3^. However, these intensive treatments offer only modest survival benefits. Recent advances in immunotherapy, including immune checkpoint inhibitions, have demonstrated minimal clinical efficacy in PDAC ^4, 5^. Currently, anti-PD-1 therapy is approved only in the second-line setting for a rare subset of patients (˂1%) with mismatch repair deficiency ^6^.

The limited efficacy of immunotherapy in PDAC largely reflects the tumor’s low intrinsic immunogenicity and its dense, immunosuppressive microenvironment, which can constitute 80–90% of the total tumor mass ^7^. The microenvironment is shaped by its cellular composition, particularly the abundance of myeloid cells that actively suppress antitumor immunity. PDAC is increasingly recognized as a prototypical example of myeloid-driven inflammation ^8^. In late-stage mouse models, myeloid cells can comprise up to 50% of the tumor mass ^9^, while in human PDAC, the estimated ratio is one myeloid cell per 2.1 tumor cells ^10^. Among these populations, tumor-associated macrophages (TAMs) are key mediators of resistance to chemotherapy ^11, 12^, radiotherapy ^13^, and immune checkpoint inhibition ^14^. Collectively, these findings underscore the myeloid compartment as a therapeutic vulnerability in PDAC. However, key challenges remain in effectively targeting myeloid cells in PDAC. A deeper understanding of pancreatic cancer biology, particularly myeloid cell function, is needed for developing more effective therapies and improving patient outcomes.

Beyond immunosuppression, myeloid cells actively remodel the PDAC TME through diverse signaling pathways. Among these signals, WNT ligands—secreted glycoproteins essential for pancreatic growth and differentiation—have emerged as context-dependent regulators of immune and stromal dynamics ^15^. Notably, myeloid cells are a major source of WNTs during tissue injury ^16^ and tumorigenesis ^17, 18^, yet their contribution in PDAC remains poorly defined.

Here, we demonstrate that noncanonical WNT ligands are predominantly produced by myeloid cells—particularly monocytes and macrophages—in human PDAC tissues and KPC mouse models. We further demonstrate that WNT-producing myeloid cells exhibit substantial functional heterogeneity. Hexb⁺ tissue-resident macrophages, which progressively accumulate to coomprise ∼50% of the immune compartment in advanced PDAC and serve as the dominant, stable source of noncanonical WNTs that sustain immunosuppressive myeloid states. In contrast, recruited TREM2⁺ monocyte-derived macrophages respond to WNT signaling by upregulating ARG1, directly suppressing CD8⁺ T-cell activation. Myeloid-specific blockade of WNT secretion increases iNOS⁺ macrophages, enhances activated CD8⁺ T-cells infiltration, reduces fibroblast abundance, and limits expansion of aggressive tumor cell populations. Collectively, these findings identify a previously unrecognized immunoregulatory circuit driven by myeloid-derived noncanonical WNT signaling and demonstrate how myeloid cell origin and functional diversity converge to promote immune resistance in PDAC.

## Materials and Methods

### Mice and human tissues

Six-week-old male C57BL/6J mice and transgenic strains (Csf1r-iCre-Ai14, Csf1r-Cre-Porcn⁻/⁻, and Ms4a3-Cre-Ai14-CD68-eGFP) were used. Genotyping was performed by PCR (primer information in Supplementary Methods and Supplementary Table 8). All animal experiments were approved by the Seoul National University Hospital IACUC and conducted in an AAALAC-accredited facility. FFPE fine-needle aspiration pancreatic cancer tissues were obtained with institutional approval.

### Tumor models and cell culture

Orthotopic and subcutaneous pancreatic tumors were established using KPC-derived primary PDA cells. Tumor growth was monitored by caliper measurement and bioluminescence imaging. KPC and primary PDA cells were cultured under standard conditions. Bone marrow–derived macrophages (BMDMs) were generated and polarized or conditioned as indicated.

### Cell isolation and flow cytometry

Single-cell suspensions were prepared from pancreatic tumors by enzymatic digestion. Cells were analyzed or sorted by flow cytometry using fluorophore-conjugated antibodies (Supplementary Table 9).

### Molecular and functional assays

Gene expression was assessed by qRT–PCR, ELISA, western blotting, immunohistochemistry, immunofluorescence, and RNAscope RNA FISH. T cell proliferation assays were performed using CFSE-labeled splenic T cells treated with macrophage-conditioned media or recombinant WNT proteins. In selected experiments, arginase-1 inhibition and in vivo CD8⁺ T cell depletion were performed.

### Single-cell and spatial analyses

scRNA-seq data were processed using Cell Ranger and analyzed with Seurat, Harmony, and CellChat. TCGA-PAAD datasets were analyzed for TAM signature–based stratification. Imaging mass cytometry (IMC) was performed using metal-tagged antibodies (Supplementary Tables 10-11), and data were analyzed using the Steinbock pipeline with spatial neighborhood analysis. IMC and IF images were co-registered for integrated single-cell analysis.

### Statistics

Statistical analyses were performed using GraphPad Prism. Additional methods are provided in the supplementary material.

## Result

### TAM signature correlates with noncanonical WNT expression in human PDAC

Previous studies have reported that Wnt/β-catenin signaling is required for pancreatic carcinogenesis ^19^. However, the specific contribution of WNT ligands derived from TAM remains unclear. We analyzed transcriptomic data from 178 patients with PDAC in TCGA to examine the relationship between TAMs and WNT signaling (Supplementary Table 1). Patients were stratified into four groups—TAM-low (n=10), mid-low (n=43), mid-high (n=61), and TAM-high (n=64)—based on the expression of 37 validated TAM signature genes, including *C1QA/B/C*, *PLTP*, *CCL2*, and *FOLR2* (Figure 1A; Supplementary Figure 1A; Supplementary Table 2) ^20^. Consistent with findings in breast and endometrial cancers ^21^, a high TAM signature score was associated with poorer survival in patients with PDAC (Figure 1B).

**Figure 1.**
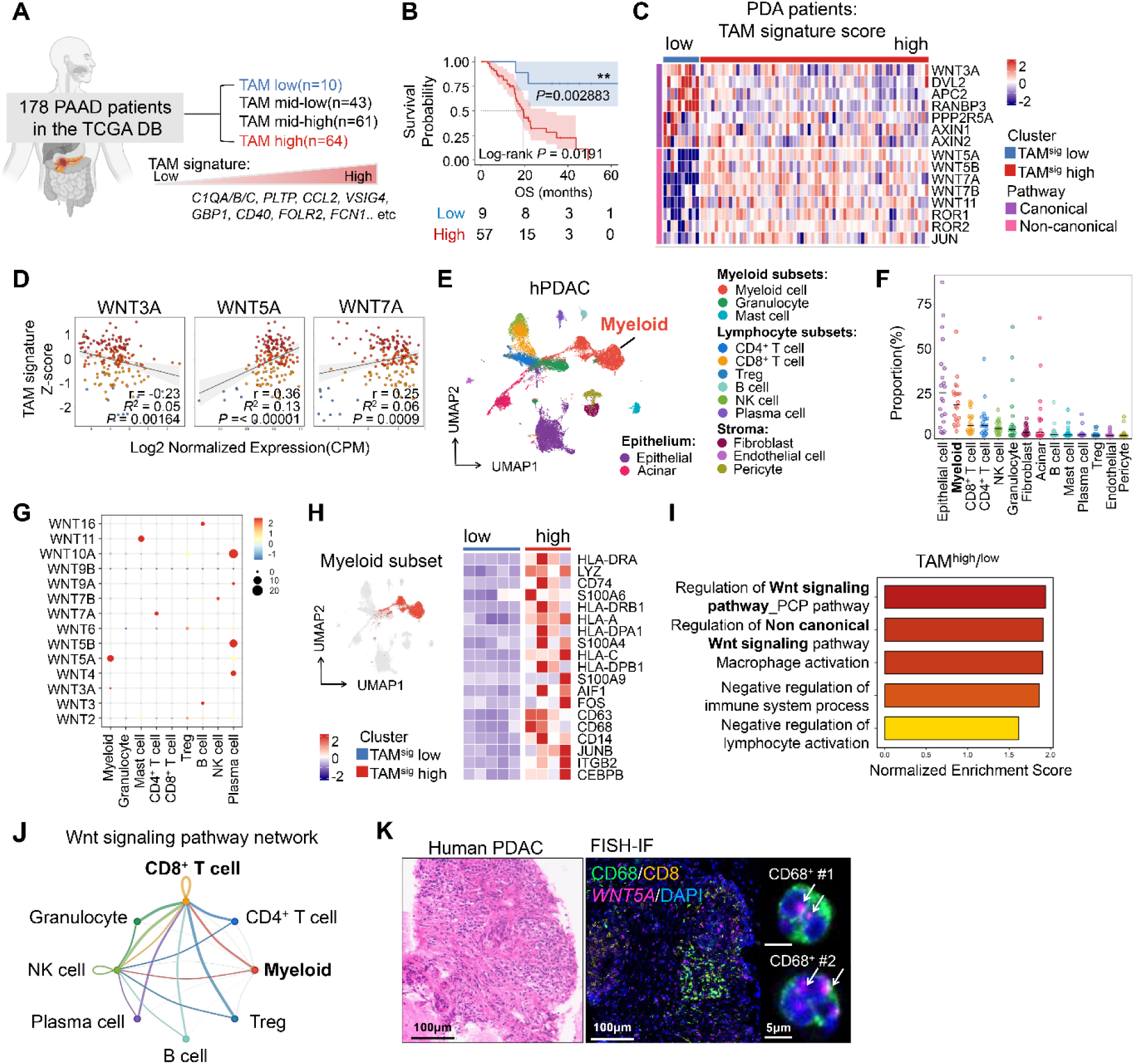
TAM abundance is associated with activation of noncanonical WNT signaling in human PDAC. (**A**) Schematic of TCGA analysis workflow. PAAD: pancreatic adenocarcinoma (**B**) Kaplan–Meier overall survival comparing TAM signature high versus low groups (shaded areas, 95% CI). The p values are calculated using a two-sided log-rank test. * *p* < 0.05; ** *p* < 0.01. (**C**) Heatmap showing the scaled expression level of representative WNT ligands and pathway components across TCGA PDAC patients stratified by TAM signature score. **(D**) Correlation between TAM signature score and log_2_-normalized expression (CPM) of WNT3A, WNT5A, and WNT7A across TCGA PDAC tumors (n=178). (**E**) Uniform Manifold Approximation and Projection (UMAP) plot of a public human PDAC scRNA-seq dataset, showing that the integrated cell map consists of 12 annotated cell types (n=53,740). (**F**) Per-sample cell-type composition from the scRNA-seq cohort showing the proportion of each annotated population across tumors. (**G**) Dot plot showing the expression of WNT ligand gene markers in each cell type. (**H**) UMAP showing myeloid cell clusters from the human PDAC scRNA-seq dataset (left). Heatmap of scaled expression of TAM marker genes across myeloid clusters used to stratify patients into TAM-low, TAM-mid, and TAM-high based on the TAM signature score. (**I**) Gene set enrichment analysis (GSEA) comparing TAM high versus low PDA patients. (**J**) Circle plot displaying putative WNT signaling pathway interactions inferred from human scRNA-seq data, with edge width indicating the inferred strength of communication between cell types. (**K**) Representative image of H&E and *WNT5A* RNAscope in situ hybridization with CD68 (green) with Nuclei (blue) in human PDAC tissue. Arrows indicate *WNT5A* RNA (pink) signals.

We evaluated the relationship between TAM infiltration and WNT signaling by analyzing the expression of 15 canonical and noncanonical WNT pathway signature genes in PDAC (Figure 1C) ^22^. Canonical WNT pathway genes (*WNT3A*, *DVL2, APC2, RANBP3, PPP2R5A, AXIN1,* and *AXIN2*) were significantly enriched in TAM-low tumors, whereas TAM-high tumors exhibited elevated expression of noncanonical WNT-associated genes (*WNT5A, WNT5B, WNT7A, WNT7B, WNT11, ROR1, ROR2,* and *JUN*). Scatter plot analysis revealed a negative correlation between *WNT3A* expression and the TAM signature score, while *WNT5A* and *WNT7A* showed positive correlations (Figure 1D). High expression of noncanonical *WNT7A (HR=3.27, log-rank P=0.0006)* and *WNT7B* (*HR=2.56, log-rank P=0.0135*) was positively correlated with worse survival outcomes (Supplementary Figure 1B), supporting a link between TAM infiltration and noncanonical WNT signaling in PDAC.

We next identified the cellular sources of WNT ligands within the pancreatic TME. Publicly available single-cell RNA sequencing data (GSE155698) from 17 human PDAC tumors and three normal pancreas samples were reanalyzed ^23^. Among 14 distinct cell types identified by UMAP analysis, myeloid cells, including TAMs, represented the second most abundant population after epithelial cells, comprising ∼19.33% of the TME (Figure 1E–F; Supplementary Figure 1C–D; Supplementary Table 3). Analysis of WNT ligand expression across immune cell populations showed that myeloid cells are the dominant source of noncanonical WNT ligands, particularly *WNT5A* (Figure 1G).

We investigated how TAM infiltration intensity influences transcriptomic features by stratifying the dataset into TAM-high (n=4), TAM-mid (n=8), and TAM-low (n=5) groups according to the TAM signature score (Figure 1H) ^24^ and performed gene set enrichment analysis. TAM-high tumors were significantly enriched for noncanonical WNT signaling pathways, including the planar cell polarity pathway, as well as pathways associated with macrophage activation and suppression of lymphocyte activation (Figure 1I). CellChat analysis showed extensive WNT signaling interactions between myeloid and CD8^+^ T cells, with WNT ligands predominantly secreted by myeloid cells (Figure 1J). We further validated that *WNT5A* transcripts colocalize with CD68^+^ macrophages in PDAC biopsy samples (Figure 1K).

Collectively, these findings indicate an association between TAM infiltration and activation of noncanonical WNT pathways in PDAC.

### Macrophage-derived WNT signaling sustains PDAC growth and epithelial plasticity

We generated macrophage-specific *Porcupine* (*Porcn)* knockout mice to investigate the role of macrophage-derived WNT in pancreatic cancer directly. Porcn, an endoplasmic reticulum–resident O-acyltransferase ^25^ required for WNT palmitoylation and secretion, was deleted by crossing *Porcn^fl/fl^* mice with *Csf1r-Cre* mice, generating *Csf1r-Cre;Porcn^fl/fl^*animals and abolishing WNT secretion from Csf1r⁺ myeloid cells (Figure 2A; Supplementary Figure 2A–F).

**Figure 2.**
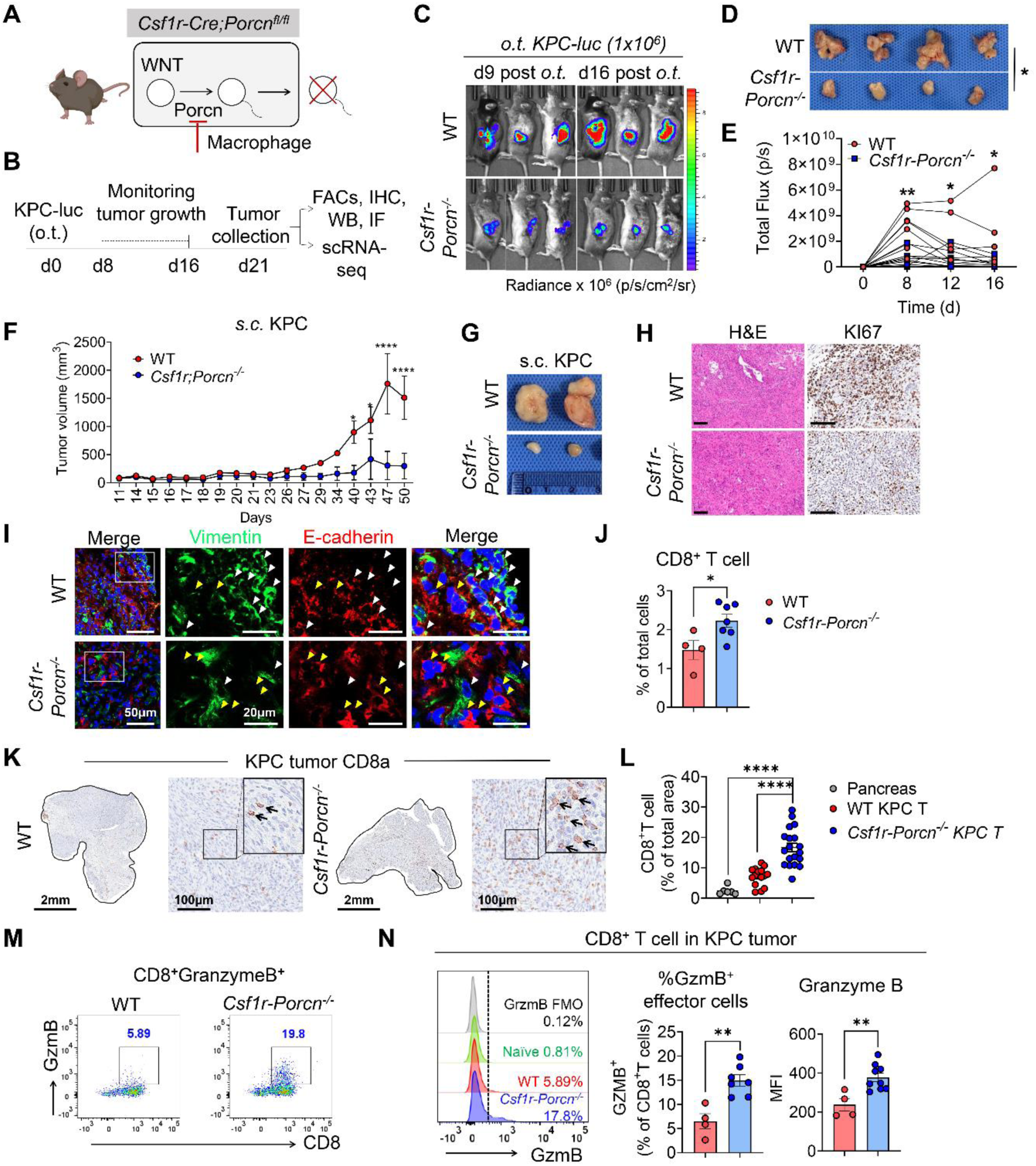
Macrophage-derived WNT secretion promotes PDAC growth and immune exclusion. (**A**) Schematic of myeloid lineage restricted ablation of WNT secretion by deleting Porcn in Csf1r-expressing myeloid cells. (**B**) Experimental timeline for orthotopic implantation of KPC-luc cells. (**C**) Representative IVIS images of orthotopic KPC-luc tumors in WT and *Csf1r-Porcn^-/-^*mice at the indicated days post-implantation. (**D**) Representative gross images of orthotopic pancreatic tumors at day 21. (**E**) Quantification of total bioluminescence flux over time for orthotopic tumors (n=8 WT, n=9 *Csf1r-Porcn*^-/-^). (**F**) Tumor growth kinetics in the subcutaneous KPC model (n=3 WT, n=3 *Csf1r-Porcn*^-/-^). (**G**) Representative gross images of subcutaneous tumors at day 50. (**H**) Representative images of H&E and Ki67 immunohistochemistry (IHC) staining of orthotopic tumors from WT and *Csf1r-Porcn^-/-^* mice. Scale bars, 100 μm. (**I**) Representative IF staining for vimentin (green) and E-cadherin (red) with nuclear counterstain (DAPI, blue). Arrowheads denote Vimentin^hi^ (yellow arrowheads) and E-cadherin^hi^ (white arrowheads) areas/cells. (**J**) Flow cytometry quantification of CD8^+^ T cells (% of total cells) in orthotopic tumors. (**K**) Representative whole-section CD8α IHC of orthotopic tumors with magnified insets (n=3 biologically independent samples). (**L**) Quantification of CD8^+^ T cell area in normal pancreas and orthotopic tumors. Each dot represents an independent imaging field/section. (**M**) Representative flow cytometry plots of granzyme B (GzmB) expression in tumor-infiltrating CD8^+^ T cells from WT and *Csf1r-Porcn^-/-^* tumors. (**N**) Representative histogram overlays (including FMO/naïve pancreas controls as indicated) and quantification of % GzmB^+^ CD8^+^ effector cells and GzmB mean fluorescence intensity (MFI). The data are shown as means +/± S.E.M. For (C,D,E,F), Two-way ANOVA test. For (J, N), Two-tailed unpaired t-test, n=5 WT, n=9 Csf1r-Porcn-/-. For (L), one-way ANOVA * *p* <0.05, ** *p* <0.01, *** *p* <0.001, **** *p* <0.0001.

In the KPC (Kras^G12D/+^; Trp53^R172H/+^; Pdx1-Cre) orthotopic pancreatic tumor model, Csf1r expression was largely restricted to macrophages and monocytes, validating this approach for studying myeloid-derived WNT signaling during pancreatic cancer progression (Supplementary Figure 2G–L). The orthotopic KPC model reproduced key pathological features of PDAC, including pronounced desmoplasia with increased collagen deposition and marked macrophage infiltration compared to the normal pancreas at 21 days post-implantation (Supplementary Figure 2M–N).

Ablation of macrophage-derived WNT secretion significantly impaired KPC tumor growth in both orthotopic (Figure 2B–E) and subcutaneous models (Figure 2F–G). Consistent with impaired tumor growth, *Csf1r-Porcn^-/-^* tumors showed reduced epithelial proliferation, as indicated by decreased Ki67 staining (Figure 2H). Immunofluorescence analysis revealed altered epithelial marker expression, with WT tumors exhibiting increased E-cadherin^lo^Vimentin^hi^ cells compared with *Csf1r-Porcn^-/-^* tumors (Figure 2I), indicating that macrophage-derived WNT signaling within the PDAC microenvironment promotes EMT transition.

### Macrophage-derived WNT sustains an immunosuppressive tumor microenvironment

We next examined how macrophage-derived WNT influences the immune landscape of KPC tumors. WT KPC tumors exhibited a “cold” immune microenvironment characterized by low CD8^+^ T cell infiltration. In contrast, *Csf1r-Porcn^-/-^* mice showed markedly increased CD8^+^ T cell infiltration, as confirmed by flow cytometry and immunohistochemistry (Figure 2H, 2J-L). Tumor-infiltrating CD8^+^ T cells exhibited enhanced cytotoxic activity, with significantly elevated granzyme B frequency and mean fluorescence intensity (Figure 2M-N). These findings demonstrate that macrophage-derived WNT is essential for sustaining an immunosuppressive PDAC microenvironment that limits infiltration of granzyme B^+^ cytotoxic CD8^+^ T cells and supports tumor growth.

### Loss of myeloid-derived WNTs reduces proliferative basal-like tumor cell populations while expanding CD8⁺ T cells, CD4⁺ T cells, and NK cells in PDAC

We performed scRNA-seq on tumors isolated from WT and *Csf1r-Porcn^-/-^* mice to define the landscape of TME shaped by myeloid-derived WNTs (Figure 3A). We identified 11 distinct clusters annotated using canonical gene markers, functional cell type signatures, and differentially expressed genes (DEGs) (Supplementary Figure 3A-B, Figure 3B, and Supplementary Table 4).

**Figure 3.**
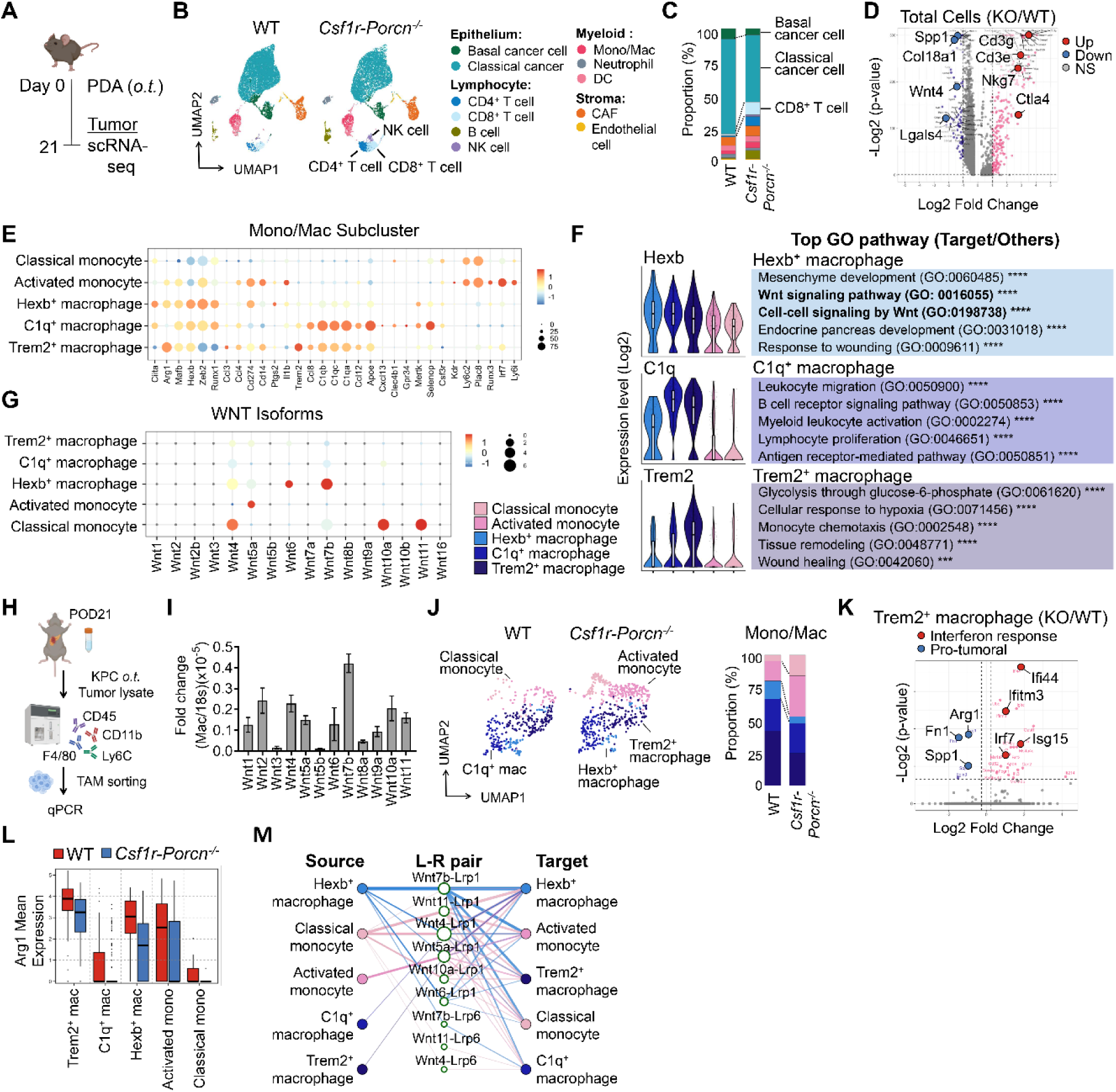
Single-cell RNA-sequencing reveals HEXB⁺ macrophages as a dominant source of noncanonical WNT signaling in PDAC. (**A**) Experimental schematic for mouse PDAC single-cell RNA-sequencing (scRNA-seq). (**B**) UMAP plots displaying the integrated cell map from WT and *Csf1r-Porcn^-/-^* tumors, which consists of 11 annotated cell types (WT cell count=11,986, KO cell count=11,301). (**C**) Stacked bar plots showing the proportion of total cell-type compartments in WT versus *Csf1r-Porcn^-/-^* tumors. (**D**) Volcano plot depicting differentially expressed genes between *Csf1r-Porcn^-/-^* and WT tumors. Red dots represent genes expressed at higher levels in *Csf1r-Porcn^-/-^* tumors, whereas blue dots represent genes expressed at higher levels in WT tumors. The y axis denotes −log_2_(*p* value) values, and the x axis shows log_2_ fold-change values. (**E**) Dot plot showing marker expression and frequency across monocyte/macrophage subclusters. (**F**) Violin plots (left) display the representative expression pattern across different subtypes of macrophage. Gene Ontology (GO) enrichment analysis (right) based on DEGs of macrophage subsets. Enrichment significance is shown as adjusted P values. *** *p* <0.001, **** *p* <0.0001. (**G**) Dot plot showing expression patterns of WNT isoforms across monocyte/macrophage subclusters. (**H**) Schematic of tumor-associated macrophage (TAM) isolation from orthotopic KPC tumors (CD45^+^CD11b^+^F4/80^+^Ly6C^-^gating) for qPCR-based validation of Wnt transcripts. (**I**) Representative graph showing mRNA levels of Wnt isoforms in TAMs from orthotopic KPC tumors. (**J**) UMAP (left) of re-clustered monocyte/macrophage populations comparing WT and *Csf1r-Porcn^-/-^*tumors, with accompanying stacked bar plot (right) showing shifts in monocyte/macrophage subcluster proportions. (**K**) Volcano plot of differentially expressed genes in TREM2^+^ macrophages comparing *Csf1r-Porcn^-/-^*versus WT tumors. Two-sided Wilcoxon rank-sum test. (**L**) Arg1 expression across monocyte/macrophage subclusters in WT and *Csf1r-Porcn^-/-^*tumors. (**M**) CellChat-based inference of putative ligand–receptor communication (WNT signaling) across mono/mac subset.

Among the epithelial populations, *Krt19*^+^ pancreatic cancer cells segregated into two clusters corresponding to the Moffitt subtypes ^26^. Consistent with tumor growth data, *Csf1r-Porcn^-/-^*tumors exhibited reduced *Gata6*^+^ classical and *S100a2*^+^ *Mki67^+^Top2a^+^* proliferating basal cancer cells, indicating that myeloid-derived WNT signals support cycling basal-like cell expansion (Supplementary Figure 3C). The most significantly expanded clusters in *Csf1r-Porcn^-/-^* tumors were adaptive immune populations (Figure 3C). Specifically, CD8^+^ T cells increased from 1.0% to 8.6%, CD4^+^ T cells from 0.9% to 7.1%, and NK cells from 0.2% to 1.3%, consistent with flow cytometry results (Figure 3C and Figure 2J). Transcriptomic profiles supported these findings, with increased *Cd3e*, *Cd3g*, and *Nkg7* expression in *Csf1r-Porcn^-/-^* tumors, consistent with enhanced T and NK cells infiltration or activation (Figure 3D). Conversely, genes associated with tumor progression, including *Spp1* ^27^, *Col18a1* ^28^, and *Lgals4* ^29^, were downregulated, a less tumor-permissive microenvironment following myeloid-derived WNT loss.

### Hexb⁺ tissue-resident macrophages represent a dominant WNT-producing macrophage state in PDAC

Unsupervised clustering of the Cd68⁺ monocyte-macrophage compartment identified five major myeloid subtypes (Figure 3E). Two Ly6c2⁺Plac8⁺ classical monocyte populations were distinguished, one exhibiting elevated *Irf7* and *Il1b* expression consistent with an activated inflammatory phenotype. Gene ontology (GO) analysis showed enrichment of type I interferon–associated pathways, including “response to interferon-β” and “pattern recognition receptor signaling,” supporting an immune-stimulatory role for this subset (Supplementary Figure 3D).

TAMs, which comprised most myeloid cells, segregated into Hexb⁺, C1q⁺, and Trem2⁺ populations (Figure 3E). GO analysis indicated that C1q⁺ macrophages were enriched for antigen receptor–mediated signaling and lymphocyte proliferation, consistent with previous reports ^11^. In contrast, Trem2⁺ macrophages exhibited transcriptional programs associated with glycolytic metabolism and adaptation to tissue hypoxia (Figure 3F).

Hexb⁺ macrophages showed significant enrichment of WNT signaling genes, including *Tcf7l2, Bcl9, Ror1, Macf1*, and *Taz*, supporting a specialized role in regulating mesenchymal developmental programs through WNT ligand production as tissue-resident macrophages (TRMs) (Supplementary Table 5) ^30^. Consistent with this role, Hexb⁺ macrophages preferentially expressed the WNT ligands *Wnt4* and *Wnt7b* and noncanonical WNT receptors, including *Ror1* and *Ryk*, indicating capacity for noncanonical WNT signaling (Figure 3F–G; Supplementary Figure 3E–3F). Among other subsets, classical monocytes were the second most prominent WNT source, primarily expressing *Wnt6*. In vivo analysis of sorted CD45⁺CD11b⁺Ly6G⁻Ly6C⁻ TAMs from KPC tumors confirmed elevated *Wnt7b* expression, supporting enrichment of noncanonical WNT programs in the PDAC myeloid compartment (Figure 3H–I).

Comparison of myeloid subcluster proportions between WT and *Csf1r-Porcn⁻/⁻* tumors revealed a marked reduction in Hexb⁺ macrophages (14% to 5%), and a concurrent expansion of activated monocytes (15% to 31%) (Figure 3J). The most pronounced transcriptional changes occurred in Trem2⁺ macrophages (Figure 3K). In *Csf1r-Porcn⁻/⁻* tumors, Trem2⁺ macrophages—previously characterized as immunosuppressive ^31^ —upregulated interferon-responsive genes (*Ifi44, Ifitm3*, and *Irf7*) while downregulating protumoral genes (*Arg1, Fn1*, *Spp1*). Although Trem2⁺ macrophages exhibited the highest *Arg1* expression, all five myeloid populations expressed *Arg1* to varying degrees in a partially WNT-dependent manner (Figure 3L).

WNT-focused ligand–receptor interaction analysis identified Hexb⁺ macrophages as the principal WNT “source,” with predicted autocrine signaling and paracrine signaling to activated monocytes and Trem2⁺ macrophages (Figure 3M). Predicted interactions were dominated by noncanonical WNT ligands produced by Hexb⁺ macrophages (Wnt4, Wnt7b, Wnt5a) engaging Lrp1/Lrp6 receptors on recipient myeloid cells, supporting a model in which WNT-producing resident TAMs establish local signaling circuits that reinforce WNT-responsive states and promote downstream suppressive programs, including Arg1 expression in Trem2⁺ macrophages.

### Spatial lineage-resolved IMC defines the ontogeny and functional specialization of pancreatic tumor–associated macrophages

We established a lineage-resolved spatial imaging strategy combining genetic fate mapping with high-dimensional IMC to overcome the limitations of dissociative single-cell approaches in resolving macrophage ontogeny and spatial organization. We used Ms4a3-Cre;Ai14^fl/fl^;hCD68-EGFP double-reporter mice, in which monocyte-derived cells irreversibly express tdTomato ^32^ and macrophages express EGFP, enabling unambiguous in situ discrimination between monocyte-derived macrophages (tdTomato⁺EGFP⁺) and TRMs (tdTomato⁻EGFP⁺) (Supplementary Figure 4A).

KPC cells were orthotopically implanted into the pancreas of double-reporter mice, after which tumors were harvested for spatial immune profiling by IMC using a 16-marker panel (Supplementary Table 10). Regions of interest (ROIs) were selected from immunofluorescence images to capture tumor-adjacent pancreas, compact lesions, and late expansile tumors, thereby enabling spatial and temporal analysis of macrophage dynamics during PDAC progression (Figure 4A–B). Following co-registration of immunofluorescence and IMC nuclear images, all ROIs were retained for single-cell segmentation and quantitative spatial analysis (Supplementary Figure 4B–C).

**Figure 4.**
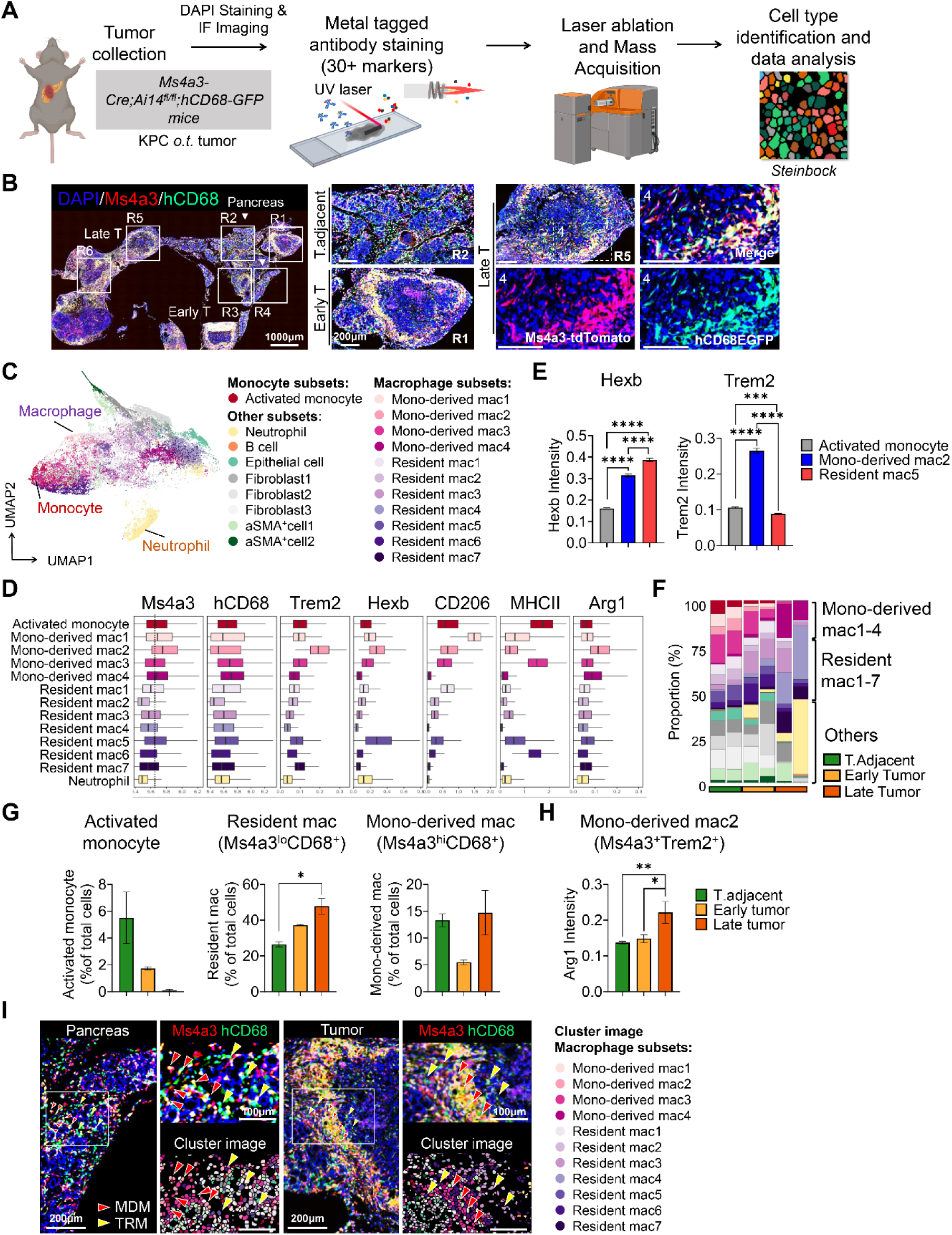
Integrated fate-mapping and imaging mass cytometry define the ontogeny and spatial organization of tumor-associated macrophages in PDAC. **(A)** Schematic of the experimental workflow. Ms4a3Cre-Ai14^TdTomato^/hCD68-EGFP double-reporter mice bearing orthotopic KPC tumors were processed for imaging mass cytometry (IMC), staining with a metal-tagged antibody panel (16 markers). (**B**) Representative fluorescence images from orthotopic KPC tumors illustrating regions of interest (ROIs) selected to capture tumor-adjacent pancreas and temporally distinct tumor regions (early/compact lesions versus late/expansile lesions). (**C**) UMAP visualization of IMC single-cell data, which consists of 20 annotated cell types (Cell count=29,913). (**D**) Box plot showing the expression of markers across monocyte/macrophage subclusters. (**E**) Quantification of mean intensity in macrophage subtypes. (**F**) Relative abundance of monocyte-derived and resident macrophage subclusters in tumor-adjacent (n=2), early tumor (n=2), and late tumor (n=2), displayed as stacked proportions. (**G**) Proportional quantification of activated monocytes, resident macrophages (Tdtomato^lo^EGFP^+^), and monocyte-derived macrophages (Tdtomato^+^EGFP^+^) across tumor-adjacent, early, and late tumor regions. (**H**) ARG1 signal intensity in the monocyte-derived mac2 state (Tdtomato^+^EGFP^+^Trem2^+^) across tumor-adjacent, early, and late tumor regions. (**I**) Representative images illustrating the spatial distribution of resident versus monocyte-derived macrophage programs. IF image showing Ms4a3 (red) reporter and hCD68 (green) labeling; meta-cluster visualization summarizing segmentation-based classification of monocyte-derived (red arrowheads) versus resident macrophages (yellow arrowheads). The data are shown as means +/± S.E.M. For (E, H), One-way ANOVA test * *p* <0.05, ** *p* <0.01, *** *p* <0.001, **** *p* <0.0001.

Across all ROIs, 29,913 cells were classified into 21 distinct cell types using Phenograph clustering, including activated monocytes, multiple TAM subsets, and non-myeloid stromal compartments (Figure 4C, Supplementary Figure 4D–E, Supplementary Table 6). Ms4a3 expression was confined to activated monocytes and monocyte-derived macrophages and was absent from tissue-resident macrophages and non-myeloid populations, validating in situ fate assignment (Supplementary Figure 4F). Hexb expression was highest in Ms4a3⁻CD68⁺ tissue-resident macrophages, whereas Trem2 was selectively enriched in Ms4a3⁺CD68⁺ monocyte-derived macrophages (Figure 4D–E), consistent with macrophage states defined by scRNA-seq.

Spatial quantification revealed marked differences in macrophage dynamics during tumor progression that were not fully captured by single-cell transcriptomic analyses. Activated monocytes were enriched in the tumor-adjacent pancreas but were less abundant in early lesions, indicating that monocyte recruitment is not a dominant feature of early PDAC formation. Tissue-resident macrophages (Ms4a3^lo/−^CD68⁺) progressively accumulated from tumor-adjacent regions through early and late tumors, constituting the dominant expanding macrophage population during PDAC progression (Figure 4F–G). Monocyte-derived macrophages (Ms4a3⁺CD68⁺) remained less abundant throughout tumor development.

Despite their lower abundance, monocyte-derived macrophages underwent functional reprogramming within advanced tumors. The Trem2⁺ monocyte-derived macrophage subset Mac2 exhibited significantly increased Arg1 expression in late tumors (Figure 4H), mirroring transcriptional programs identified by scRNA-seq. Spatial mapping demonstrated that Trem2⁺Arg1⁺ monocyte-derived macrophages were intermingled with Hexb⁺ tissue-resident macrophages across normal pancreas and tumor regions (Figure 4I), indicating functional coupling rather than spatial segregation.

Collectively, these lineage-resolved spatial analyses reveal a hierarchical organization of the PDAC macrophage compartment, in which tissue-resident macrophages progressively accumulate to establish a WNT-rich niche, while less abundant Trem2⁺ monocyte-derived macrophages acquire Arg1-dependent immunosuppressive function within this niche. This integrated genetic and spatial imaging approach robustly validates our single-cell findings and reveals a previously unrecognized, ontogeny-dependent organization of macrophage-mediated immune suppression in PDAC.

### Myeloid WNT signaling organizes a CD8⁺ T cell–restricted tumor architecture in PDAC

High-dimensional 35-plex IMC analysis was performed on orthotopic KPC tumors from WT and *Csf1r-Porcn⁻/⁻* mice to determine how myeloid-derived WNT signaling shapes the spatial organization of the PDAC microenvironment. Unsupervised phenotyping delineated major epithelial, stromal, and immune compartments, encompassing multiple epithelial states and myeloid subsets (Figure 5A, Supplementary Figure 5A, Supplementary Table 7).

**Figure 5.**
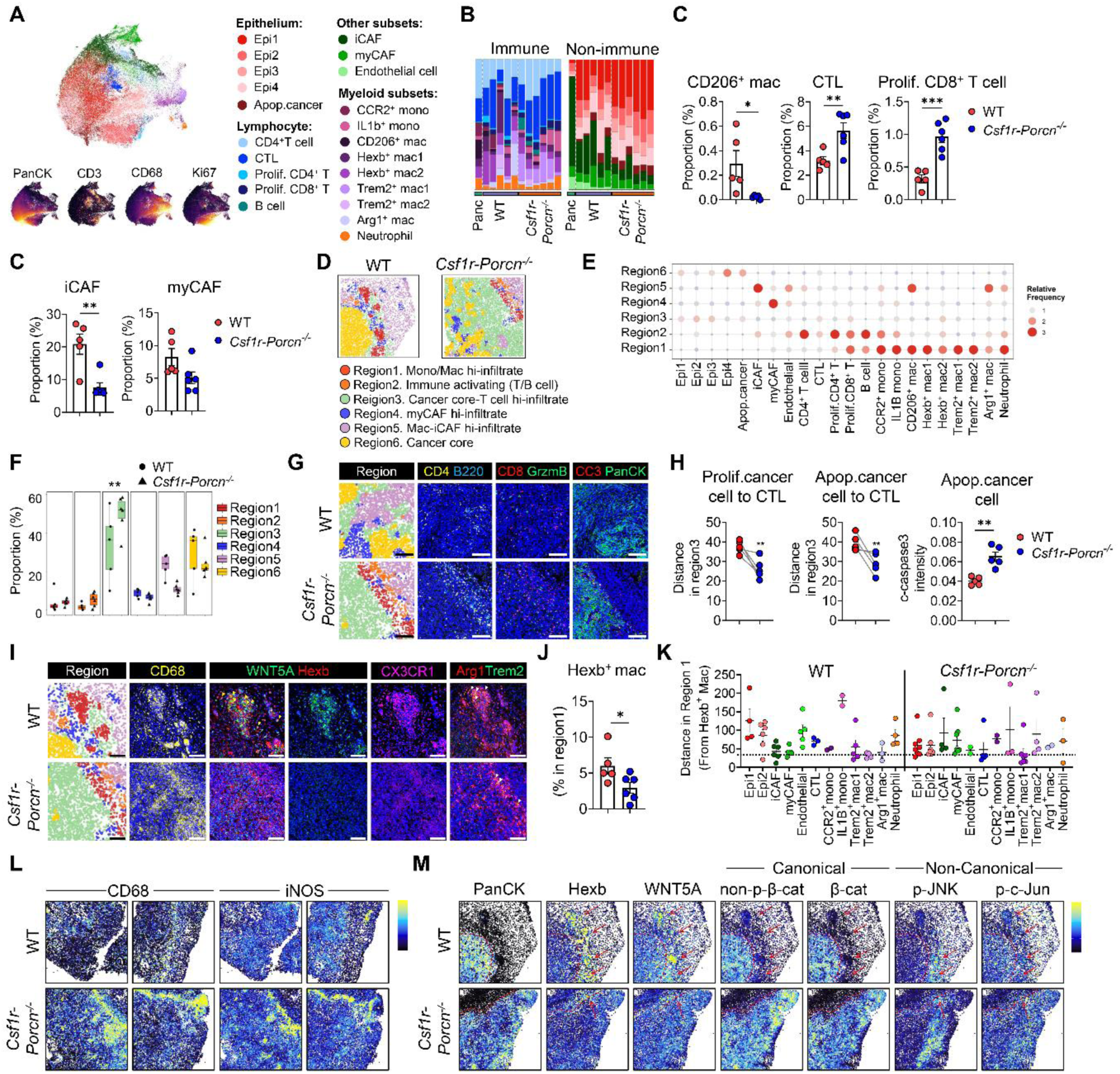
Myeloid-derived WNT5A establishes a HEXB⁺ macrophage niche that spatially restricts cytotoxic immune access in PDAC. (**A**) UMAP visualization of single-cell IMC data from pancreas, WT, and *Csf1r-Porcn^-/-^* orthotopic KPC tumors showing major epithelial, stromal populations, and immune subsets. Marker-intensity feature plots (bottom) show representative lineage markers for PanCK, CD3, CD68, and KI67. (**B**) Stacked bar plots summarizing the relative abundance of immune (left) and non-immune (right) compartments across normal pancreas and tumors from WT and *Csf1r-Porcn^-/-^* mice. (**C**) Quantification of CD206⁺ macrophages, cytotoxic CD8⁺ T cells (CTL), proliferating CD8⁺ T cells, and iCAF, myCAF in *Csf1r-Porcn⁻/⁻* tumors compared with WT. (**D**) LisaClust-based spatial domain mapping of representative WT and *Csf1r-Porcn^-/-^*tumor ROIs. (**E**) Dot plot showing region-enriched cell-type compositions (relative frequency) across the six LISA regions, including immune subsets, epithelial states, and CAF subtypes. (**F**) Distribution of region area fractions in WT versus *Csf1r-Porcn^-/-^* tumors across all analyzed ROIs. (**G**) Left, the LISA-cluster annotated spatial domain map is directly aligned with the IMC channel images (right). Representative IMC images of CD4, B220, CD8, Granzyme B, PanCK, and cleaved caspase-3 (CC3) in WT and *Csf1r-Porcn^-/-^* tumors. (**H**) Quantification of CTL distance to proliferating and apoptotic cancer cells in Region 3, and CC3 intensity in apoptotic cancer cells. (**I**) Representative macrophage-niche visualization in WT and *Csf1r-Porcn^-/-^* tumor ROI. Left, the LISA-cluster annotated spatial domain map is directly aligned with the IMC channel images (right), confirming that TREM2^+^ and HEXB^+^ macrophages correspond to their respective marker-positive signals in the raw IMC channels. Right, IMC channels show CD68, WNT5A (overlaid with HEXB), CX3CR1, ARG1 (overlaid with TREM2). Yellow arrows indicate WNT5A^+^ HEXB^+^ macrophages within these niches. (**J**) Percentage of HEXB⁺ macrophages among CD68⁺ cells within Region 1. (**K**) Nearest-neighbor distances between Hexb⁺ macrophages and others. (L) Marker overlay images for CD68 and iNOS expressions. (**M**) Marker expression in representative ROIs showing PanCK, HEXB, and WNT5A, together with canonical readouts (non–phospho–β-catenin, β-catenin) and noncanonical readouts (p-JNK, p-c-JUN). Dotted lines delineate PanCK^+^ tumor nests. Outlined areas indicate HEXB/WNT5A-enriched niches with preferential enrichment of noncanonical signaling activity. The data are shown as means +/± S.E.M. Each dot represents one ROI. * *p* <0.05, ** *p* <0.01, *** *p* <0.001, **** *p* <0.0001.

Compositional analysis revealed broad remodeling of the TME following myeloid-specific *Porcn* deletion, characterized by a reduction in CD206⁺ macrophages and a concurrent expansion of cytotoxic lymphocytes and cancer-associated fibroblast (CAF) subsets (Figure 5B–C). These compositional changes were next evaluated for their association with altered spatial tissue organization, using lisaClust to partition tumors into spatially conserved microenvironments. This analysis identified six distinct regions with characteristic cellular compositions and uncovered marked differences in the spatial organization of these regions between WT and *Csf1r-Porcn⁻/⁻* tumors (Figure 5D–F, Supplementary Figure 5B). Representative IMC ROIs are shown in Supplementary Figure 6A-C.

Annotation demonstrated that WT tumors were dominated by macrophage- and CAF-enriched regions (regions 4 and 5) and cancer cell–enriched regions (region 6), indicative of immune-exclusion. Conversely, *Csf1r-Porcn⁻/⁻* tumors exhibited expanded immune-enriched regions with lymphocyte accumulation (regions 2 and 3) (Figure 5F, Supplementary Figure 5C).

Spatial proximity analysis indicated that myeloid–epithelial and macrophage–CAF interfaces were more compact in WT tumors. Specifically, Ccr2⁺ were significantly closer to EMT-like epithelial niches, and Hexb⁺ macrophages were nearer to iCAF and myCAF subsets in WT compared to *Csf1r-Porcn⁻/⁻* tumors (Supplementary Figure 5D–G). This spatial uncoupling in knockout tumors was associated with reduced fibroblast abundance and collagen deposition, supporting a role for myeloid WNT signaling in maintaining CAF–TAM interaction niches (Supplementary Figure 5H–I).

Consistent with this spatial architecture, CTLs in WT tumors were largely excluded from PanCK⁺ tumor nests and confined to peripheral regions, whereas *Csf1r-Porcn⁻/⁻* tumors displayed increased CD8⁺ T cell infiltration into tumor regions (Figure 5G, Supplementary Figure 6C). Spatial distance analysis confirmed that CTLs in *Csf1r-Porcn⁻/⁻*tumors were positioned significantly closer to both proliferating and apoptotic tumor cells, accompanied by increased cleaved caspase-3 intensity, indicative of enhanced tumor cell apoptosis upon disruption of myeloid WNT signaling (Figure 5H).

### Hexb⁺ macrophages define a WNT5A noncanonical signaling niche in PDAC

Within WT tumors, Hexb⁺ macrophages were preferentially enriched in a distinct macrophage-dense region (region 1), which coincided with elevated WNT5A signal, defining a discrete WNT-producing niche that was markedly diminished in *Csf1r-Porcn⁻/⁻* tumors (Figure 5I and J). Trem2⁺ macrophages expressing Arg1 were frequently positioned adjacent to Hexb⁺ macrophages within this region in WT tumors (Figure 5I and 5K, Supplementary Figure 5H), further suggesting spatial coupling between a WNT-producing resident-like macrophage population and an Arg1-associated suppressive macrophage state. Finally, in *Csf1r-Porcn⁻/⁻* tumors, CD68⁺ macrophages formed a dense, continuous rim encircling tumor regions, along which iNOS expression was strongly and selectively induced (Figure 5L). This spatially organized inflammatory macrophage band in region 1 was not observed in WT tumors, indicating a WNT-dependent switch from suppressive TAM niches to an inflammatory barrier-like macrophage organization.

To directly assess WNT pathway activity in situ, we examined spatial co-localization of signaling markers. In WT tumors, WNT5A signal overlapped with Hexb-enriched macrophage regions and was accompanied by localized activation of noncanonical WNT downstream effectors, as indicated by increased p-JNK and p-cJUN signals in adjacent macrophages. In contrast, *Csf1r-Porcn⁻/⁻* tumors exhibited markedly attenuated p-JNK and p-cJUN activation and loss of spatial concordance with Hexb⁺ macrophage regions (Figure 5M). Canonical β-catenin–associated readouts were not similarly enriched, supporting preferential engagement of noncanonical WNT signaling within this niche.

Together, these spatial analyses demonstrate that Hexb⁺ macrophages establish a WNT5A-driven, noncanonical signaling niche that organizes suppressive macrophage states and enforces immune exclusion in PDAC.

### Noncanonical WNT5A–JNK signaling programs ARG1 expression in tumor-educated macrophages

Given our earlier observation that CD8⁺ T cell infiltration was increased in *Csf1r-Porcn⁻/⁻* KPC tumors, we next asked whether tumor-educated macrophages suppress CD8⁺ T cell proliferative capacity through WNT. To address this, WT or *Csf1r-Porcn⁻/⁻* bone marrow–derived macrophages (BMDMs) were first conditioned with KPC tumor cell culture supernatant, after which macrophage-conditioned media (hereafter referred to as TAM-CM) were collected and applied to CFSE-labeled CD8⁺ T cells stimulated with anti-CD3/CD28 (Figure 6A). CD8⁺ T cells cultured with TAM-CM derived from WT macrophages exhibited significantly reduced proliferation compared with those cultured with TAM-CM from Porcn-deficient macrophages (Figure 6B–C).

**Figure 6.**
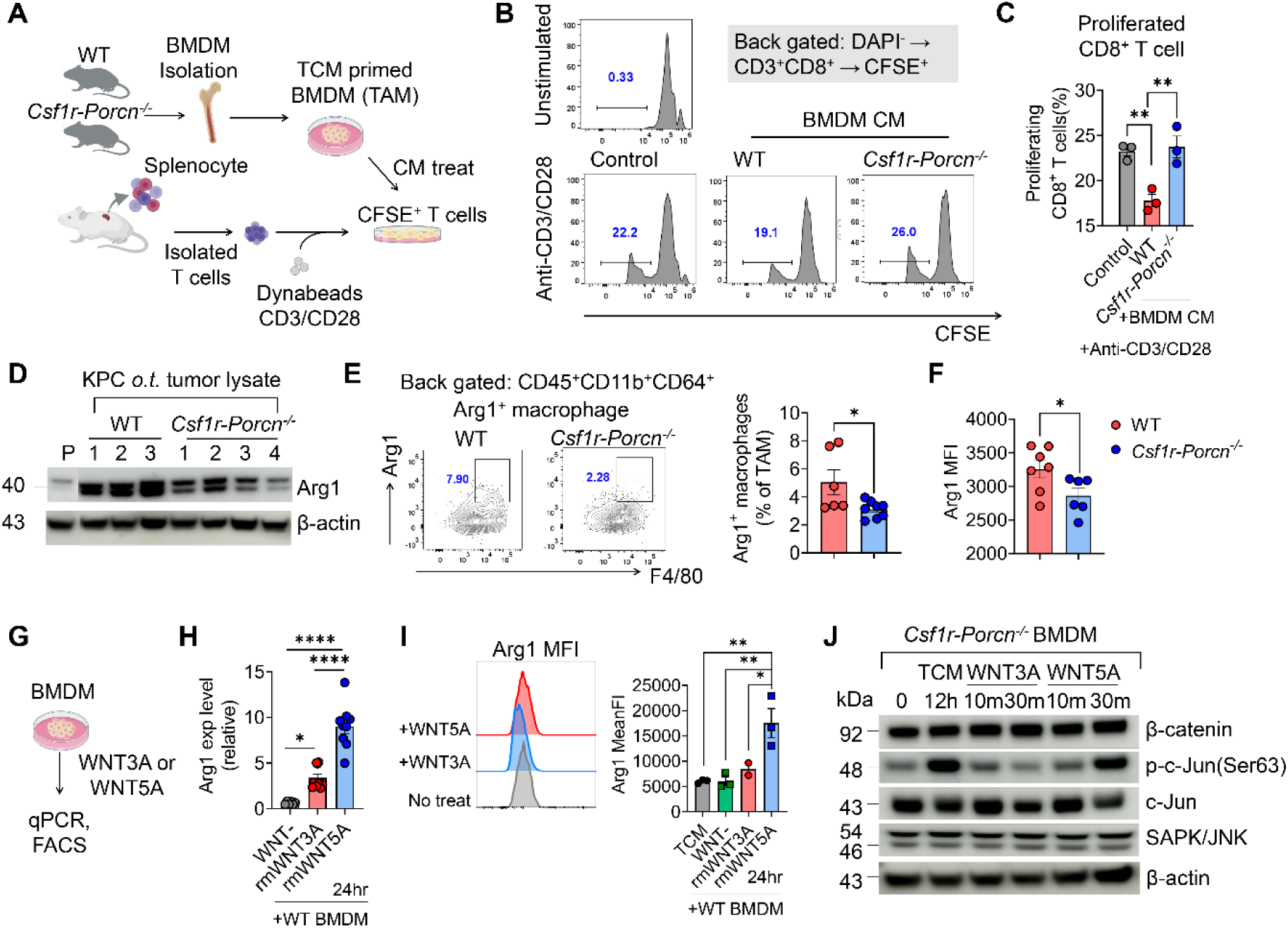
Noncanonical WNT5A–JNK/c-Jun signaling in macrophages drives ARG1-mediated suppression of CD8⁺ T cell proliferation. (**A**) Schematic of the in vitro T cell proliferation assay. Bone marrow–derived macrophages (BMDMs) from WT or *Csf1r-Porcn^-/-^*mice were primed with tumor-conditioned medium (TCM) to generate TAM-like macrophages, and CM was transferred to CFSE-labeled primary CD8⁺ T cells activated with anti-CD3/CD28 Dynabeads. (**B**) Representative CFSE dilution histograms of activated CD8⁺ T cells cultured with control medium (no CM) or CM derived from WT or *Csf1r-Porcn^-/-^*TAMs. Cells were gated as DAPI^-^CD3^+^CD8^+^ CFSE^+^. (**C**) Quantification of proliferating CD8⁺ T cells under the indicated CM conditions. Each dot represents an independent biological replicate. (**D**) Western blot of ARG1 in lysates from orthotopic KPC tumors harvested from WT and *Csf1r-Porcn^-/-^*mice (WT n=3, KO n=4 tumors). β-actin, loading control. (**E**) Representative flow cytometry plots showing ARG1 expression (left) and Arg1^+^ macrophage frequency (right) in tumor macrophages gated as CD45^+^CD11b^+^CD64^+^ from WT and *Csf1r-Porcn^-/-^*KPC tumors. (**F**) Quantification of ARG1 mean fluorescence intensity (MFI) in TAMs (CD45^+^CD11b^+^CD64^+^) from WT and *Csf1r-Porcn^-/-^* tumors. (**G**) Schematic of recombinant WNT stimulation in macrophages. WT BMDMs were treated with recombinant mouse WNT3A (canonical) or WNT5A (noncanonical) and analyzed for Arg1 induction by qPCR and flow cytometry. (**H**) qPCR quantification of Arg1 induction in WT BMDMs after WNT3A or 5A stimulation (24 h), normalized to the indicated control. (**I**) Representative histograms (left) and quantification (right) of ARG1 protein MFI in WT BMDMs treated with recombinant WNT3A or 5A (24 h). (**J**) Western blot analysis of WNT pathway signaling in *Csf1r-Porcn^-/-^* BMDMs stimulated with recombinant WNT3A or 5A for the indicated times, probing β-catenin (canonical), p-c-Jun (Ser63), c-Jun, and SAPK/JNK (noncanonical related); β-actin, loading control. The data are shown as means +/± S.E.M. For (C,H,I), One-way ANOVA test, For (E,F), Two-tailed unpaired t-test, * *p* <0.05, ** *p* <0.01, *** *p* <0.001, **** *p* <0.0001.

We next examined whether this difference was associated with altered Arg1 expression downstream of macrophage WNT signaling. Arg1 protein levels were markedly reduced in whole KPC tumor lysates from *Csf1r-Porcn⁻/⁻* mice compared with WT controls, whereas Arg1 expression in normal pancreas remained low (Figure 6D). Consistent with this, flow cytometric analysis demonstrated significantly decreased Arg1 expression in CD11b⁺F4/80⁺CD64⁺ tumor macrophages from *Csf1r-Porcn⁻/⁻* tumors (Figure 6E–F).

To define the upstream WNT ligand and signaling pathway regulating Arg1 expression, BMDMs were treated with recombinant canonical mouse WNT3A or noncanonical WNT5A. WNT5A robustly induced Arg1 expression at both transcript and protein levels, whereas WNT3A had minimal effect (Figure 6G–I). Consistent with activation of noncanonical WNT signaling, WNT5A treatment induced phosphorylation of SAPK/JNK and c-Jun without altering β-catenin levels in macrophages (Figure 6J).

### Macrophage-derived WNT–ARG1 signaling limits CD8⁺ T cell–dependent antitumor immunity in vivo

To determine whether the suppressive effect CD8^+^ T cell proliferation was mediated by Arg1, we exposed CD8⁺ T cells to TAM CM supplemented with CB-1158 (Numidargistat)^33^, a selective small-molecule arginase-1 inhibitor (Figure 7A). CB-1158 treatment fully restored CD8⁺ T cell proliferation to levels comparable to those cultured with *Porcn*-deficient TAM CM, indicating that the proliferation defect of CD8^+^ T cells was largely dependent on Arg1 activity (Figure 7B-C).

**Figure 7.**
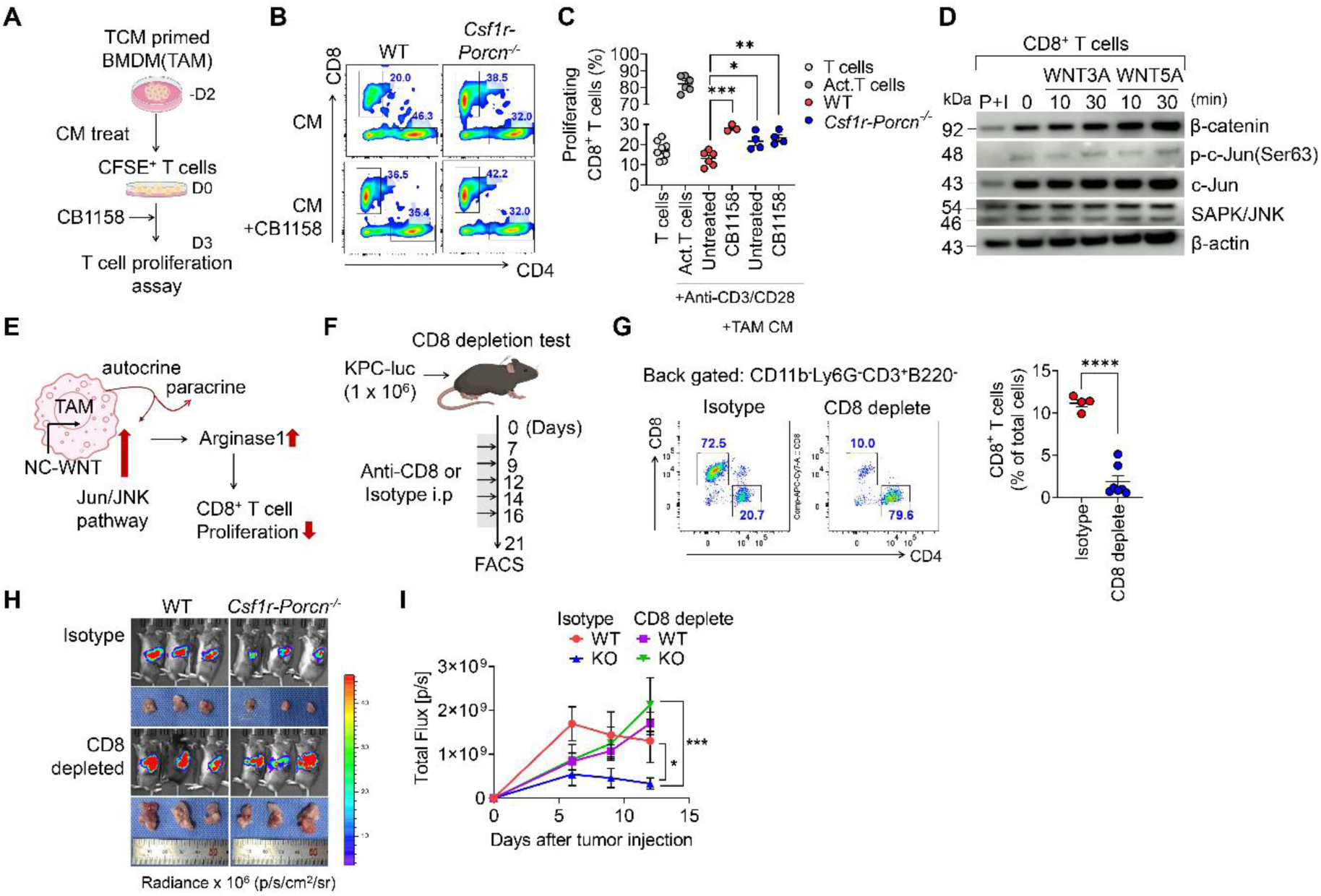
Macrophage-derived WNT–ARG1 signaling suppresses CD8⁺ T cell proliferation and constrains antitumor immunity in vivo. (**A**) Schematic of the in vitro T cell proliferation assay. KPC TCM primed BMDMs were generated from WT or *Csf1r-Porcn^-/-^* mice, and CM was collected. CFSE-labeled CD8^+^ T cells were stimulated with anti-CD3/CD28 and cultured with TAM CM ± CB-1158 (Numidargistat) followed by assessment of T cell proliferation by CFSE dilution. (**B**) Representative flow cytometry plots showing proportion of CD8^+^ T cells cultured with WT or *Csf1r-Porcn^-/-^* TAM CM in the presence or absence of CB-1158. (**C**) Quantification of proliferating CD8^+^ T cells (% of total cells) under the indicated conditions. (**D**) Western blot analysis of purified CD8^+^ T cells treated with recombinant WNT3A or 5A for the indicated times, probing β-catenin and noncanonical pathway (p-c-Jun, c-Jun, SAPK/JNK); β-actin served as a loading control. P+I, PMA + Ionomycin. (**E**) Schematic summarizing an autocrine noncanonical WNT-JNK/c-Jun program in TAMs that induces ARG1 and suppresses CD8^+^ T cell proliferation. (**F**) Schematic of in vivo CD8^+^ T cell depletion in the orthotopic KPC model. Mice were implanted with KPC-luc cells (1×10^6^) and intraperitoneal injection with anti-CD8α antibody or isotype control according to the indicated schedule, followed by immune profiling and tumor assessment. (**G**) Representative flow cytometry plots and quantification confirming efficient depletion of intratumoral CD8^+^ T cells in anti-CD8α-treated mice (back-gated on CD11b^-^Ly6G^-^CD3^+^B220^-^cells). (**H**) Representative IVIS imaging of orthotopic tumors from WT and *Csf1r-Porcn^-/-^* mice treated with isotype or anti-CD8α antibody. (**I**) Quantification of total bioluminescence flux over time (n=6 isotype, n=7 CD8 depleted). The data are shown as means +/± S.E.M. For (C), One-way ANOVA test, For (G), Two-tailed unpaired t-test, n=4 Isotype, n=7 CD8 depleted. For (I), Two-way ANOVA test. * *p* <0.05, ** *p* <0.01, *** *p* <0.001, **** *p* <0.000

We next asked whether WNT ligands exert a direct signaling effect on CD8⁺ T cells. To address this, purified CD8⁺ T cells were treated with recombinant WNT3A or WNT5A, and activation of downstream signaling pathways was assessed by immunoblotting. Neither WNT3A nor WNT5A induced robust activation of canonical or noncanonical WNT signaling in CD8⁺ T cells, as evidenced by minimal changes in β-catenin levels and only marginal phosphorylation of SAPK/JNK and c-Jun (Figure 7D). These results indicate that WNT ligands alone are insufficient to directly suppress CD8⁺ T cells and support an indirect, macrophage-mediated mechanism underlying WNT-dependent suppression of CD8⁺ T cell proliferation (Figure 7E).

Finally, to determine whether the pro-tumoral role of myeloid-derived WNT signaling is mediated through CD8^+^ T cells in vivo, we performed CD8 depletion in the orthotopic KPC tumor model (Figure 7F). Mice were treated with anti-CD8α antibody or isotype control following tumor implantation, and efficient depletion of CD8^+^ T cells was confirmed by flow cytometry (Figure 7G). Strikingly, CD8^+^ T cell depletion largely abrogated the tumor-growth defect observed in *Csf1r-Porcn^-/-^* mice, leading to increased tumor burden to levels comparable to WT controls (Figure 7H–I). These results indicate that the restraint of tumor outgrowth upon loss of myeloid WNT secretion is predominantly CD8^+^ T cell–dependent, consistent with a macrophage WNT–ARG1 axis that limits CD8^+^ T cell proliferative capacity and antitumor immunity.

## Discussion

High numbers of tumor-associated macrophages (TAMs) in PDAC correlate with poor clinical outcomes, making macrophages an attractive therapeutic target ^34, 35^. However, prior strategies aimed at broadly targeting the myeloid compartment have been limited by compensatory expansion of immunosuppressive myeloid populations and loss of macrophage immunostimulatory functions ^12^. Notably, high TAM density at the tumor–stroma interface is associated with improved chemotherapy responsiveness, underscoring the context-dependent tumoricidal roles of macrophages ^36^. In this study, we show that selective disruption of macrophage-derived WNT secretion, without depleting macrophages, results in a striking restriction of PDAC growth, with tumor volumes reduced by six- to seven-fold in an aggressive KPC model (Figure 1D, 1G). This growth restraint is accompanied by the emergence of a dense, iNOS-enriched macrophage rim (Figure 5M) and is largely dependent on CD8⁺ T cell activity (Figure 7H). These observations support therapeutic strategies that selectively disrupt pro-tumoral myeloid signaling architectures, such as macrophage-restricted inhibition of WNT secretion via PORCN, to reprogram the PDAC microenvironment toward effective antitumor immunity. Consistent with this concept, the OPTIMIZE phase II trial demonstrated clinical benefit of mitazalimab, a CD40 agonistic antibody that reprograms myeloid cells, combined with mFOLFIRINOX in metastatic pancreatic cancer ^37^.

This study provides the first evidence of macrophage-associated noncanonical WNT5A in human PDAC biopsies using combined in situ FISH and immunofluorescence (Figure 1K). Consistent with this observation, reanalysis of published human PDAC scRNA-seq datasets revealed detectable *WNT5A* transcripts within myeloid cell populations (Figure 1G). Notably, transcriptomic analyses further demonstrated a striking enrichment of noncanonical WNT/PCP signaling pathways in TAM-high PDAC tissues (Figure 1I). In contrast to the prevailing focus on tumor cell intrinsic, canonical β-catenin–driven WNT signaling in cancer ^38, 39^, our data reposition tumor-associated macrophages as a central and previously overlooked source of noncanonical WNT activity in human PDAC.

Across species, the dominant macrophage-derived WNT ligands appear to be context dependent. In human PDAC tissues, *WNT5A* was readily detected at the transcript level within macrophage-enriched regions (Figure 1K), whereas mouse macrophages exhibited more heterogeneous WNT ligand expression at the transcript level, with Hexb⁺ resident macrophages preferentially expressing *Wnt4* and *Wnt7b* in scRNA-seq analyses. Notably, despite this transcriptional heterogeneity, WNT5A protein was strongly enriched within tumor tissues in mice, as revealed by spatial IMC analysis (Figure 5I), and mouse BMDMs expressed Wnt5a in vitro (Supplementary Figure 2E). Together, these observations suggest that WNT5A represents a functionally dominant noncanonical WNT ligand within the PDAC microenvironment, whose biological impact is more accurately captured at the protein and transcript level than by dissociative transcriptomic approaches alone.

Importantly, this macrophage-enriched, noncanonical WNT landscape was spatially segregated from canonical WNT activity. Although KPC tumors lack RNF43 mutations, present in up to ∼10% of human PDAC ^40^, active β-catenin was selectively enriched within tumor epithelial cores (Figure 5M). In contrast, stromal regions displayed low canonical WNT activity but were characterized by prominent noncanonical features, including abundant WNT5A expression. This spatial compartmentalization suggests that canonical and noncanonical WNT programs operate in distinct cellular and anatomical domains within PDAC. In this context, early clinical experience with WNT974, a first-in-class PORCN inhibitor that demonstrated tolerability in phase I studies ^41^, may need to be reinterpreted not solely through tumor-intrinsic WNT dependence, but also through its potential effects on stromal and myeloid WNT signaling niches.

WNT-mediated immune suppression is not a novel concept in cancer biology. Previous studies have shown that canonical WNT signaling endows tumor-infiltrating CD4⁺ T cells with immunosuppressive properties in both human and mouse pancreatic ductal adenocarcinoma ^42^. WNT signaling has also been implicated in immune evasion in melanoma by impairing dendritic cell–dependent T cell cross-priming via the activity of indoleamine 2,3-dioxygenase-1 ^43, 44^. It has been shown to regulate the recruitment and function of multiple immune populations, including myeloid-derived suppressor cells, NK cells, and regulatory T cell ^45^. What our study newly reveals is a macrophage-driven organization of WNT biology in PDAC, in which myeloid-derived noncanonical WNT signaling governs CD8⁺ T cell exclusion and proliferative restraint (Figure 6). While c-Jun phosphorylation (Figure 6J) has been implicated in the establishment of immunosuppressive, pro-tumorigenic macrophage phenotypes ^46^, the precise molecular mechanisms by which noncanonical WNT signaling directly induces Arg1 expression in macrophages remain to be elucidated.

One of the most conceptually informative aspects of this study is the lineage-resolved spatial dissection of macrophage populations in PDAC achieved through the combination of macrophage fate–mapping double-reporter mice and integrated IF/IMC analysis. This approach enabled precise, in situ quantification of embryonically derived TRMs and monocyte-derived macrophages, overcoming a major limitation of dissociative single-cell methodologies. Consistent with prior lineage-tracing studies using Flt3-Cre–based reporter systems ^47^, Ms4a3⁻CD68⁺ TRMs exhibited low expression of CD11b and MHCII, confirming their embryonic origin. In the normal pancreas, resident and monocyte-derived macrophages were present at comparable proportions (Figure 4G). In contrast, PDAC progression was marked by a progressive and selective expansion of TRMs, which constituted up to ∼50% of all cells in advanced tumors. Importantly, this increase reflected a specific enrichment of functionally distinct subsets rather than a uniform expansion of resident macrophages. The Hexb⁺ subset emerged as the dominant myeloid population in late-stage tumors (Figure 4F) and was identified as the principal source of WNT ligands within the TME (Figure 3G). Notably, *Hexb* has been defined as a core microglial gene ^48^, consistent with the embryonic origin and tissue-resident identity of pancreatic resident macrophages. Consistent with observations by Zhu et al. ^47^, we found that pancreatic tissue-resident macrophages were closely associated with fibrotic features of PDAC (Supplementary Figure 5F-5I). This finding is clinically relevant, as the dense fibrotic stroma and profound immunosuppression that characterize PDAC are key contributors to its limited response to current therapies. In many cancer types, monocyte-derived macrophages constitute the major myeloid population ^49^. However, PDAC represents a notable exception, with tissue-resident macrophages emerging as the dominant macrophage subset. This distinction has important implications for the design of macrophage-targeted therapies in PDAC.

In summary, our study provides new insight into macrophage biology in PDAC by demonstrating that lineage identity is a critical determinant of macrophage function. While lineage-dependent macrophage heterogeneity has only recently begun to be explored in cancer contexts, our findings identify embryonically derived tissue-resident macrophages as a dominant source of noncanonical WNT signals that organize suppressive myeloid programs and reinforce CD8⁺ T cell exclusion in PDAC. This work shifts the focus from macrophage abundance to lineage- and signal-specific function, clarifying why indiscriminate myeloid targeting has yielded limited benefit and identifying WNT-dependent macrophage programs as tractable therapeutic vulnerabilities in PDAC.

## Abbreviations used in this paper

APC: [antigen presenting cell](https://www.gastrojournal.org/action/doSearch?type=quicksearch&text1=antigen%2Bpresenting%2Bcell&field1=AllField)
Arg1: [arginase-1](https://www.gastrojournal.org/action/doSearch?type=quicksearch&text1=arg1&field1=AllField)
BMDM: bone marrow-derived macrophage
CAF: cancer-associated fibroblast
CB1158: numidargistat
CD&#x0023;: [cluster of differentiation](https://www.gastrojournal.org/action/doSearch?type=quicksearch&text1=cluster%2Bof%2Bdifferentiation&field1=AllField)
CM: [conditioned medium](https://www.gastrojournal.org/action/doSearch?type=quicksearch&text1=conditioned%2Bmedium&field1=AllField)
CTL: [cytotoxic T lymphocyte](https://www.gastrojournal.org/action/doSearch?type=quicksearch&text1=cytotoxic%2BT%2Blymphocyte&field1=AllField)
DC: [dendritic cell](https://www.gastrojournal.org/action/doSearch?type=quicksearch&text1=dendritic%2Bcell&field1=AllField)
ECM: [extracellular matrix](https://www.gastrojournal.org/action/doSearch?type=quicksearch&text1=extracellular%2Bmatrix&field1=AllField)
EMT: epithelial-mesenchymal transition
IMC: [imaging mass cytometry](https://www.gastrojournal.org/action/doSearch?type=quicksearch&text1=imaging%2Bmass%2Bcytometry&field1=AllField)
JNK: c-Jun N-terminal kinase
PAAD: [pancreatic adenocarcinoma](https://www.gastrojournal.org/action/doSearch?type=quicksearch&text1=pancreatic%2Badenocarcinoma&field1=AllField)
PDAC: [pancreatic ductal adenocarcinoma](https://www.gastrojournal.org/action/doSearch?type=quicksearch&text1=pancreatic%2Bductal%2Badenocarcinoma&field1=AllField)
ROI: region of interest
ROR1: receptor tyrosine kinase-like orphan receptor 1
ROR2: [receptor tyrosine kinase-like orphan receptor 2](https://www.gastrojournal.org/action/doSearch?type=quicksearch&text1=ror2&field1=AllField)
scRNA-seq: [single cell RNA sequencing](https://www.gastrojournal.org/action/doSearch?type=quicksearch&text1=single%2Bcell%2BRNA%2Bsequencing&field1=AllField)
TAM: [tumor associated macrophage](https://www.gastrojournal.org/action/doSearch?type=quicksearch&text1=tumor%2Bassociated%2Bmacrophage&field1=AllField)
TCGA: [The Cancer Genome Atlas](https://www.gastrojournal.org/action/doSearch?type=quicksearch&text1=The%2BCancer%2BGenome%2BAtlas&field1=AllField)
TCM: tumor-conditioned medium
TME: tumor-microenvironment
TRM: [tissue resident macrophage](https://www.gastrojournal.org/action/doSearch?type=quicksearch&text1=tissue%2Bresident%2Bmacrophage&field1=AllField)
WNT: [WNT ligand](https://www.gastrojournal.org/action/doSearch?type=quicksearch&text1=WNT%2Bligand&field1=AllField)
WT: wild-type

## Acknowledgments

We thank members of our Translational Immunology Laboratory for helpful discussions and suggestions. The gag/pol/env and luciferase vectors were kindly provided by Prof. Hye-won Youn (Department of Nuclear Medicine, Seoul National University College of Medicine).

## CRediT Authorship Contributions

Na Hyun Kim, MS (Conceptualization: Equal; Formal analysis: Lead; Investigation: Lead; Methodology: Lead; Visualization: Lead; Writing-original draft: Lead; Writing-review & editing: Lead)

Young Min Song, MS (Formal analysis: Supporting; Visualization: Supporting, Software: Lead)

San Sung Kwon, MS (Formal analysis: Supporting; Visualization: Supporting, Software: Lead)

Sang Hyub Lee, PhD (Resources: Supporting)

Eun Na Kim, PhD (Formal analysis: Supporting)

Jung Joo Hong, PhD (Resources: Supporting)

Seung Hyeok Seok, PhD (Conceptualization: Equal; Supervision: Lead; Funding acquisition: Lead)

Yi Rang Na, PhD (Conceptualization: Lead; Supervision: Lead; Formal analysis: Lead; Funding acquisition: Lead; Writing-review & editing: Lead)

