## Supplementary material for "A macrophage-derived noncanonical WNT niche drives immune exclusion in pancreatic cancer"

### Supplementary Information

#### Supplementary Materials and Methods

##### Mice

Six-week-old male C57BL/6J mice were purchased from Koatech (Gyeonggi-do, South Korea). *Csf1r*$^{iCre}$*;Ai14*$^{fl/fl}$ mice were generated by crossing *Ai14*$^{fl/fl}$ (B6.Cg-*Gt(ROSA)26Sor*$^{tm14(CAG\text{-}tdTomato)Hze}$/J, Jax stock number #007914)[1] mice with *Csf1r-iCre* (FVB-Tg (Csf1r-cre/Esr1*) 1Jwp/J, Jax stock number #019098)[2]. *Csf1r*$^{Cre}$*;Porcn*$^{fl/fl}$ mice were generated by crossing *Porcn*$^{fl/fl}$ (129S-*Porcn*$^{tm1.1Vdv}$/J, Jax stock number #020994)[3] mice with *Csf1r-Cre* (C57BL/6-Tg (Csf1r-cre) 1Mnz/J, Jax stock number #029206)[4]. *Ms4a3*$^{Cre}$*;Ai14*$^{fl/fl}$*;CD68*-eGFP double-reporter mice were generated by crossing *Ms4a3-Cre* mice (C57BL/6J-*Ms4a3*$^{em2(cre)Fgnx}$/J, Jax stock number #036382)[5] with *Ai14*$^{fl/fl}$ reporter mice (B6.Cg-*Gt(ROSA)26Sor*$^{tm14(CAG\text{-}tdTomato)Hze}$/J, Jax stock number #007914)[1] and *hCD68*-eGFP reporter mice (C57BL/6-Tg(CD68-EGFP)1Drg/J, Jax stock number #026827)[6]. Genotypes of all transgenic mice were distinguished by PCR using the primer sets (Supplementary Table 8). All animal experiments were approved by the Institutional Animal Care and Use Committee of Seoul National University Hospital (SNUH-IACUC; accession number SNU-240415-3), and animals were maintained in the facility accredited by AAALAC International (#001169) in accordance with the Guide for the Care and Use of Laboratory Animals (8th ed., National Research Council, 2010).

##### Human PDAC tissues

Fine-needle aspiration paraffin-embedded sections from patients with pancreatic cancer tissues were provided by Professor Sang-Hyub Lee (Department of Gastroenterology, Seoul National University Hospital, Seoul, South Korea). The specimen represented an invasive pancreatic body cancer and was fixed in 10% Neutral Buffered Formalin for 5 hours before paraffin embedding.

##### Establishment of pancreatic cancer mouse models

Male mice aged 7-8 weeks were anaesthetized via intraperitoneal injection of ketamine (10 mg/kg) combined with xylazine. Orthotopic pancreatic injections were conducted as previously described [7]. After shaving the left abdominal flank, the peritoneal cavity was accessed through an incision in the left upper quadrant. For the establishment of the orthotopic pancreatic cancer model, $10^6$ primary mouse pancreatic ductal adenocarcinoma (PDA) cells derived from the KPC tumor were resuspended in 20 µl suspension of 50% Matrigel in phosphate-buffered saline (PBS) and slowly injected into the pancreatic tail. For the subcutaneous model, 100 µl of a KPC cell suspension containing $10^6$ cells was injected into the right flank of each mouse to generate syngeneic tumors. Tumor growth was monitored twice weekly using caliper measurements. Tumor volume ($T_V$) was calculated using the formula: $T_V = 0.5 \times L \times W^2$, where $L$ represents tumor length, and $W$ represents tumor width. In vivo bioluminescence imaging was performed using an IVIS Imaging System (PerkinElmer, MA, USA) after intraperitoneal administration of D-luciferin (150 mg/kg body weight). Mice were anesthetized with isoflurane and imaged 10 minutes after D-luciferin injection. Average radiance, expressed as photons per second per cm$^2$ per steradian (p/s/cm2/sr), was quantified within a circular region of interest using Living Image software (PerkinElmer).

##### Cell lines and cell culture

KPC (*Kras*$^{G12D/+}$*;Trp53*$^{R172H/+}$*;Pdx1-Cre;Luc2*) cell lines were obtained from Jennifer Morton and Saadia Karim (Ximbio, catalog no. 153474). Luciferase expression was introduced into KPC cells by retroviral transduction to enable bioluminescence imaging. Primary mouse PDA tumor cells were established from orthotopic KPC tumors harvested three weeks after tumor induction. Dissected tumor-bearing pancreata were minced and enzymatically dissociated using collagenase V (Sigma-Aldrich, C9263) and DNase I (Sigma-Aldrich, D5025), followed by filtration and cancer cells enrichment before culture (see "Cell isolation from tissues"). The KPC cell line and primary mouse PDA cells were cultured in high-glucose DMEM (Welgene) supplemented with 10% fetal bovine serum (FBS; Gibco), 1% penicillin/streptomycin (P/S; Gibco), and puromycin (5ug/mL) to select luciferase-expressing tumor cells. Cells were detached using 0.05% trypsin–EDTA (Gibco) and subsequently injected either orthotopically or subcutaneously into transgenic mice. KPC cells were seeded in T75 flasks to 70-80% confluence, and the conditioned medium (KPC TCM) was collected after two days.

Bone marrow-derived macrophages (BMDMs) were generated from C57BL/6 mouse bone marrow. Briefly, tibiae and femora were harvested from 6-8-week-old C57BL/6 mice. The bones were placed on ice and rinsed in a sterile dish. Under sterile conditions, the ends of each bone were transected, and the marrow cavities were flushed with culture medium using a sterile syringe. After red blood cell lysis, cells were washed and cultured in RPMI 1640 (Welgene) containing 10% FBS, 25 ng/mL macrophage colony-stimulating factor (M-CSF), 1% P/S, and 2mM L-glutamine (Gibco) for seven days to induce macrophage differentiation [8]. Mature BMDMs were detached from culture dishes using a cell scraper (SPL) and reseeded into plates according to the experimental design. BMDMs were stimulated with lipopolysaccharide (LPS; Sigma) at 100 ng/mL for 24 hours to generate inflammatory macrophages. BMDMs were treated with WNT3A (10 ng/mL) or WNT5A (500 ng/mL) for WNT signalling activation. BMDMs were cultured in KPC TCM for 24 h to generate tumor-associated macrophage (TAM)-like cells. All cells were cultured at 37°C in a humidified incubator with 5% $CO_2$. All cell lines were authenticated by short tandem repeat profiling and confirmed to be mycoplasma-free using the Myco-Sniff-Valid Mycoplasma PCR Detection Kit (MP Biomedicals).

**Cell isolation from tissues**

Mice bearing KPC tumors were euthanized three weeks after tumor induction, and tumor-bearing pancreatic tissues were dissected and finely minced using a razor blade (Razor med). Tissues were enzymatically digested in collagenase V (Sigma-Aldrich, C9263) and DNase I (Sigma-Aldrich, D5025) in PBS 5% containing FBS at 37°C for 20 minutes. Digested tissues were passed through a 70-µm cell strainer to generate single-cell suspensions, which were pelleted, and red blood cells were lysed using ACK buffer (150 mM ammonium chloride, 10 mM potassium bicarbonate, and 0.1 mM EDTA in distilled water). Cell pellets were washed and used for flow cytometry or cell sorting, or cultured to establish primary tumor cells.

**Flow cytometry and cell sorting**

Single-cell suspensions from pancreatic tumors were stained with antibodies against specific surface markers and CD16/CD32 Fc Block (Invitrogen) diluted in Brilliant Stain Buffer (BD) for 30 minutes on ice. For intracellular staining, cells were fixed and permeabilized using Fixation/Permeability Concentrate buffer (Invitrogen). After incubation, cells were washed with 1x PBS containing 5% FBS and 0.1% sodium azide. All staining procedures were performed in 96-well V-bottom plates, and all antibodies were titrated to determine optimal dilutions. Details of antibody panels, including fluorochromes and catalog numbers, are provided in Supplementary Table 9. Flow cytometry data were acquired using a FACSymphony™ system (BD) and analyzed with FlowJo™ v10.10.0 software. Tumor-infiltrating $CD11b^{+}F4/80^{+}$ TAMs were sorted using a Sony SH800S cell sorter with a 100-µm nozzle.

**Enzyme-linked immunosorbent assay (ELISA)**

ELISA-linked immunosorbent assays were performed using mouse protein Wnt5b ELISA kits (AFG Scientific) according to the manufacturer's instructions.

**Quantitative RT-PCR**

Total RNA was extracted using TriZol reagent. Reverse transcription for complementary DNA synthesis was performed using ReverTra Ace qPCR RT Master Mix (Toyobo) according to the manufacturer's instructions. Quantitative PCR reactions were prepared using TOPreal$^{TM}$ SYBR Green qPCR PreMIX (Enzynomics) and gene-specific primers. 18S rRNA was used as an internal control. Reactions were performed on a QuantStudio 5 Real-Time PCR system (Applied Biosystems). Quantitative gene expression was calculated as follows: $\Delta CT = CT_{\text{gene of interest}} - CT_{\text{internal control}}$; $\Delta\Delta CT = \Delta CT_{\text{treated group}} - \Delta CT_{\text{control group}}$; Fold change=$2^{-\Delta\Delta CT}$[9]. Primer sequences are listed in Supplementary Table 8.

**Immunostaining**

Mouse pancreatic cancer tissues were fixed in 4% paraformaldehyde (PFA; Thermo Fisher Scientific) at 4°C overnight, cryoprotected in phosphate-buffered saline (PBS) containing 30% sucrose and 0.1% sodium azide at 4°C overnight, and embedded in optimal cutting temperature (OCT) compound (Sakura). For paraffin embedding, tissues were fixed in 4% PFA at 4°C overnight, dehydration through graded ethanol, and embedded in paraffin.

For immunohistochemistry (IHC), formalin-fixed, paraffin-embedded (FFPE) tissues were sectioned at a thickness of 4μm. Sections were deparaffinized in xylene and rehydrated through graded ethanol solutions. Antigen retrieval was performed in sodium citrate buffer (pH 6.0) by heat-induced epitope retrieval for 30 minutes. Endogenous peroxidase activity was quenched with 0.3% hydrogen peroxide for 10 minutes in the dark, followed by washing in PBS. Sections were blocked with blocking buffer (Cell Signaling Technology) for 1 hour and incubated with the indicated primary antibodies overnight at 4°C. After washing with PBS, sections were incubated with horseradish (HRP)-polymer secondary antibodies (Dako) for 30 minutes, followed by signal detection using 3,3'-diaminobenzidine (DAB) substrate (Dako) and counterstaining with hematoxylin. Slides were mounted using Permount mounting medium (Thermo Fisher Scientific) diluted 1:1 with xylene.

For immunofluorescence (IF), OCT-embedded tissues were sectioned at 10 μm thickness using a cryo-microtome (Leica). Sections were blocked with IF blocking buffer (Cell Signaling Technology) for 1 hour and incubated overnight at 4°C with primary antibodies diluted 1:250 in antibody dilution buffer (Cell Signaling Technology). After washing, sections were incubated with Alexa Fluor-conjugated secondary antibodies (1:500; Invitrogen) for 1 hour at room temperature and counterstained with 4',6-diamidino-2-phenylindole (DAPI; Thermo Fisher Scientific). Slides were mounted using VECTASHIELD Plus Antifade Mounting Medium (Vector Laboratories). Details of all antibodies used are provided in Supplementary Table 9. Confocal images were acquired using a Carl Zeiss LSM 800 microscope, and whole-slide images were obtained using a Leica GT450 slide scanner. Images were processed using Fiji (ImageJ).

**Fluorescence in situ hybridization (FISH)**

WNT5A FISH was performed using the RNAscope Multiplex Fluorescent Reagent Kit (Bio-Techne ACDbio) according to the manufacturer's instructions. An RNA probe for targeting WNT5A (catalog no. 604921; Bio-Techne ACDbio) was used (Supplementary Table 9). Paraffin-embedded human pancreatic tissue slides were baked for 1 hour at 60°C and then deparaffinized. Antigen retrieval was performed for 30 minutes, followed by treatment with the protease reagent. The RNA scope probe was hybridized and amplified using amplification (AMP) reagents. Signal amplification was achieved using HRP reagent. Dual staining combining RNAscope-ISH with IF was performed according to the manufacturer's protocols (Bio-Techne ACDbio). Primary antibodies (Anti-CD68; clone KP1; Anti-CD8; clone CAL66; Abcam) were incubated overnight at 4°C. Secondary antibodies were incubated at room temperature. After washing, slides were counterstained with DAPI and mounted with VECTASHIELD Plus Antifade Mounting Medium (Vector Laboratories). Confocal images were acquired by Carl Zeiss LSM 800 and a Leica Stellaris 5 confocal microscope.

**T cell proliferation assay**

Splenocytes were isolated from the spleen of C57BL/6 mice, and T cells were enriched using the MagniSort Mouse T Cell Enrichment Kit (Invitrogen). Enriched T cells were labelled with 10 μM carboxyfluorescein succinimidyl ester (CFSE; Invitrogen) and activated for 24 hours with mouse T-Activator CD3/CD28 Dynabeads (3 μl/well; catalog no. 11456D; Gibco) in a 5% $CO_2$ incubator [10]. Purified T cells were cultured with RPMI 1640 medium (Welgene) supplemented with 10% FBS and 1% penicillin-streptomycin (P/S) in 96-well round-bottom plates (catalog no. 353077; Falcon). After activation, T cells were treated for three days with BMDM culture medium from *Csf1r-Porcn*$^{-/-}$ or wild-type (WT) mice, or with recombinant mouse WNT-3A or WNT-5A proteins (catalog no. 1324-WN and 645-WN; R&D Systems). T cell proliferation was measured using FACSymphony™ flow cytometer (BD Biosciences).

**Western Blotting**

For Western blotting, whole-cell and tumor tissue lysates were prepared in ice-cold lysis buffer supplemented with protease and phosphatase inhibitor cocktails. Lysates were mixed with 4× Laemmli Sample Buffer (Reducing; GenDEPOT, L1100-001), boiled for 10 min, and stored frozen until use. Protein concentrations were determined using the Pierce™ BCA Protein Assay Kit (Thermo Fisher Scientific, 23225), and tumor tissue lysates were adjusted to 3μg/μL prior to loading. Proteins were separated by SDS–PAGE using a Mini Gel Tank (Thermo Fisher Scientific, A25977) and transferred to membranes using Power Blotter Select Transfer Stacks (PB5310, Invitrogen). Membranes were blocked in 5% skim milk for 1 hour at room temperature with shaking, washed with PBS-T (0.05% Tween), and incubated with primary antibody diluted in 5% BSA in PBS-T overnight at 4°C. After washing, membranes were incubated with HRP-conjugated secondary antibodies. Signals were detected by SuperSignal™ West Femto Maximum Sensitivity Substrate (Thermo Fisher Scientific) and imaged using a Chemi Image system. For reprobing, membranes were stripped using Easter-Blot Western Blot Stripping Buffer (Biomax). Antibody details are provided in Supplementary Table 9.

**Arg1 inhibition assay**

WT or Csf1r-Porcn$^{-/-}$ BMDMs were treated with the arginase-1 inhibitor CB-1158 at a concentration of 10 μM for 48 hours. Following treatment, macrophage-conditioned media were collected, clarified by centrifugation, and used to stimulate $CD8^+$ T cells in subsequent T cell proliferation assays. Control conditioned media were prepared from vehicle-treated BMDMs under experimental conditions.

**In vivo mouse CD8 depletion**

For in vivo $CD8^+$ T cell depletion, mice were inoculated with $1 \times 10^6$ KPC-luc cells on day 0. $CD8^+$ T cells were depleted by intraperitoneal injection of 100 μg anti-CD8α monoclonal antibody (clone 2.43; Bio X Cell) on days 7, 9, 12, 14, and 16 after tumor implantation. Control mice received an isotypematched antibody on the same schedule. Detailed antibody information is provided in Supplementary Table 9. Mice were harvested on day 21, and the efficiency of $CD8^+$ T cell depletion and overall immune composition were assessed by flow cytometry.

**Single-cell RNA sequencing data processing and analysis**

Raw sequencing data were processed using Cell Ranger (v8.0.0) with the mm10-2020-A mouse reference genome. Libraries were prepared using the Chromium Single Cell 3' kit (v3; 10x Genomics) and sequenced on the Illumina NovaSeq 6000 platform. All downstream analyses were conducted in R (v4.3.3) using Seurat (v5.3.1) [11], Harmony (v1.2.4)[12], and clusterProfiler (v4.10.1) [13]. Quality control filtering excluded cells with fewer than 200 or greater than 8,000 detected genes or with mitochondrial gene content exceeding 20%. Gene expression data were log-normalized using a scale factor of 10,000, and the top 2,000 highly variable genes were identified using a variance-stabilizing transformation. The data were scaled, and principal component analysis was used for dimensionality reduction. Batch effects were corrected using the Harmony algorithm, with sample identity specified as the batch variable. Graph-based clustering was performed using the shared nearest neighbor algorithm at a resolution of 2.5 based on the first 10 principal components. Cell populations were visualized using UMAP projection based on Harmony-corrected embeddings. Cell types were annotated based on canonical marker gene expression. Monocyte and macrophage populations were extracted and re-independently analyzed to characterize myeloid cell heterogeneity. Following the same normalization and batch correction procedures, graph-based clustering was performed at a resolution of 0.6. One cluster exhibiting contamination signatures was excluded from downstream analysis, and the remaining subpopulations were annotated based on marker gene expression. Differential gene expression between conditions was assessed using the Wilcoxon rank-sum test implemented in the FindMarkers() function. Genes with an adjusted $P < 0.05$, expression in at least 10% of cells in either group, and absolute $\log_2$ fold change > 0.25 were considered significantly differentially expressed. Gene Ontology (GO) biological process enrichment analysis was performed using the enrichGO() function in clusterProfiler. Gene symbols were converted to Entrez identities using the org.Mm.eg.db annotation database (v3.18.0). Background gene sets were defined as all genes detected within the relevant cell population. Enrichment significance was determined using Fisher's exact test with Benjamini-Hochberg multiple-testing correction. Cell-cell communication analysis was performed using CellChat (v2.2.0)[14] with the CellChatDB.mouse ligand-receptor database. Separate CellChat objects were generated for each condition using normalized expression data. Overexpressed ligands and receptors were identified for each cell population, and communication probabilities were computed using the trimmed mean method with population-size normalization. Interactions involving fewer than 10 cells were excluded from the analysis. Pathway-level communication probabilities were calculated, and interaction networks were aggregated for downstream analysis. Network centrality scores were computed to identify dominant signaling senders and receivers. For comparative analysis between conditions, CellChat objects were merged, and differential interaction patterns were assessed to identify condition-specific signaling changes.

**TCGA-PAAD data processing and TAM signature–based stratification**

Transcriptomic and clinical data from patients with pancreatic adenocarcinoma (PAAD) were obtained from The Cancer Genome Atlas (TCGA) via the Genomic Data Commons (GDC) using the TCGAbiolinks [15, 16] R package (v2.30.4). RNA sequencing gene expression data generated using the STAR-Counts workflow were downloaded, comprising 178 primary tumor samples and four solid tissue normal samples. Raw count matrices were preprocessed by filtering lowly expressed genes using the filterByExpr() function in edgeR (v4.0.16), retaining genes with sufficient expression across samples. Library sizes were normalized using the trimmed mean of M-values method with calcNormFactors(), followed by voom transformation in limma (v3.58.1) to generate log-counts per million values with precision weights for downstream analysis. Ensembl gene identifiers were converted to HGNC symbols using biomaRt (v2.58.2), and duplicate symbols were resolved by retaining the transcript with the highest mean expression. A TAM signature was derived from a previously published 37-gene set associated with human TAM transcriptional programs (Supplementary Table 2). For each primary tumor sample, expression values of TAM signature genes were extracted from the normalized matrix and transformed into z scores by centering and scaling across samples. A TAM signature score was calculated for each tumor as the mean z score across all TAM signature genes. Patients were stratified into four groups based on TAM signature scores using the ConsensusClusterPlus algorithm (v1.66.0) with hierarchical clustering (Ward.D2 linkage), Euclidean distance, 500 bootstrap iterations, and 80% sample resampling. The optimal number of clusters (K=4) was selected based on consensus matrix stability. The four resulting clusters were labeled "Low," "Mid-Low," "Mid-High," and "High" according to ascending mean TAM signature scores. Overall survival was defined as the time from initial diagnosis to death or last follow-up. Kaplan-Meier survival curves were generated using the survival (v3.5.8) and survminer (v0.5.1) R packages, and group differences were assessed using the log-rank test. Hazard ratios were estimated using Cox proportional hazards regression models. All statistical analyses were performed in R (v4.3.3).

**Imaging mass cytometry**

Antibodies were prepared using the Maxpar X8 Conjugation Kit (Standard BioTools) according to the manufacturer's instructions. All staining steps followed the protocol provided by Fluidigm (FLDM-00073, Rev. 01). Briefly, Imaging mass cytometry (IMC) staining was performed on both FFPE and fixed-frozen tissue sections with sample-type–specific preprocessing. For FFPE sections, slides were deparaffinized in 100% xylene for 10 minutes twice, rehydrated in 100% ethanol for 10 minutes twice, and subjected to antigen retrieval before staining. Fixed-frozen sections were processed directly for staining. After preprocessing, slides were washed in Maxpar PBS (Standard BioTools), blocked with 3% BSA and 0.3% Triton X-100 in Maxpar PBS for 30 minutes at room temperature, and incubated overnight at 4 °C with a metal-conjugated antibody cocktail (Supplementary Tables 10-11) diluted in Maxpar PBS containing 0.5% BSA and 0.3% Triton X-100. Antibodies were centrifuged at 13,000g for 2 minutes to remove aggregates before use. Slides were washed twice with 0.1% Tween-20 in 1x PBS, stained with Intercalator-Ir (1:400) for 30 minutes at room temperature, rinsed in Maxpar water, air-dried for at least 20 minutes, and stored in slide boxes until acquisition. IMC data were acquired using a Hyperion Tissue Imager (Standard BioTools) with CyTOF software (v7.0.8).

**Imaging mass cytometry data analysis**

IMC data were acquired using the Hyperion Imaging System (Standard BioTools). Raw MCD files were converted into TIFF images and processed using the Steinbock pipeline[17]. Hot pixels were removed by replacing values exceeding a predefined threshold with the mean intensity of the eight surrounding pixels. Cell segmentation was performed using Mesmer, a pretrained deep learning model for nuclear and whole-cell segmentation. Single-cell expression data and spatial coordinates were extracted using the imcRtools package (v1.8.0) and compiled into a SpatialExperiment object in R (v4.3.3). Raw ion counts were transformed using an inverse hyperbolic sine (arcsinh) transformation with a cofactor of 5. Cells with an area smaller than 5 pixels were excluded as potential debris. Principal component analysis was performed on expression values using 30 components with the runPCA() function in scater (v1.30.1), followed by batch correction using the Harmony algorithm with sample identity as the batch variable. UMAP embedding was computed on Harmony-corrected principal components using the runUMAP() function. Unsupervised clustering was performed using a self-organizing map with 100 nodes, followed by consensus clustering with ConsensusClusterPlus (v1.66.0; maxK = 60, reps = 100, distance = "euclidean"). Clusters were manually annotated based on canonical marker expression patterns, and those with ambiguous profiles were excluded from downstream analysis. To further characterize myeloid cell heterogeneity, monocytes and macrophages were extracted and reclustered using FlowSOM[18] (v2.10.0), followed by consensus metaclustering. Subclusters were manually annotated based on marker expression patterns and integrated into the final cell-type annotations. Spatial relationships between cell types were analyzed using the lisaClust algorithm (v1.10.1)[19] to identify spatially colocalized cellular neighborhoods based on local cell-type composition. Pairwise cell-cell distances were calculated using the minDistToCells() function in imcRtools, and mean distances between cell-type pairs were compared between experimental groups using the Wilcoxon rank-sum test with Benjamini-Hochberg correction for multiple comparisons.

**Co-registration of IMC and IF**

IF and IMC images were aligned using the OpenCV library in Python 3.13.7. Briefly, nuclear channels from both modalities were extracted and normalized to match intensity levels. Next, IMC DNA images were scanned against IF DAPI images to identify the coordinates that maximized cross-correlation between the two modalities. At the optimal alignment score, IF images were cropped to match the IMC field of view. Co-registration was further refined using findTransformECC(), and the resulting warping matrix was applied to the remaining IF channels. Using the same cell segmentation mask, IF intensities were calculated and assigned to the corresponding IMC cells for downstream analysis. IF images for tdTomato, GFP, and DAPI were acquired using a BC43 benchtop confocal microscope.

**Statistical analysis**

Statistical analyses were performed using GraphPad Prism 8.4.3 (GraphPad software), and data are presented as mean ± SD. Student's two-tailed t-test and one or two-way ANOVA with Turkey's multiple comparison test were carried out for comparison between groups. All experiments were carried out in at least three independent replicates. Sample sizes are indicated in the figure legends. Statistical significance was defined as $P<0.05$.

**Data availability**

Human PDAC scRNA-seq datasets were obtained from the NIH Gene Expression Omnibus (GEO, https://www.ncbi.nlm.nih.gov/geo/) under accession number GSE155698 [20].

The raw scRNA-seq data generated in this study are publicly accessible on GEO under accession number GSE316305. All remaining data are provided in the article and its supplementary files.

**Code availability**

Available upon reasonable request

2. Qian BZ, Li J, Zhang H, et al. CCL2 recruits inflammatory monocytes to facilitate

breast-tumour metastasis. Nature 2011;475:222–5.

3. Liu W, Shaver TM, Balasa A, et al. Deletion of Porcn in mice leads to multiple developmental defects and models human focal dermal hypoplasia (Goltz syndrome). PLoS One 2012;7:e32331.
4. Loschko J, Rieke GJ, Schreiber HA, et al. Inducible targeting of cDCs and their subsets in vivo. J Immunol Methods 2016;434:32–8.
5. Liu Z, Gu Y, Chakarov S, et al. Fate Mapping via Ms4a3-Expression History Traces Monocyte-Derived Cells. Cell 2019;178:1509–1525.e19.
6. Iqbal AJ, McNeill E, Kapellos TS, et al. Human CD68 promoter GFP transgenic mice allow analysis of monocyte to macrophage differentiation in vivo. Blood 2014;124:e33–44.
7. Kim MP, Evans DB, Wang H, et al. Generation of orthotopic and heterotopic human pancreatic cancer xenografts in immunodeficient mice. Nature Protocols 2009;4:1670–1680.
8. Toda G, Yamauchi T, Kadowaki T, et al. Preparation and culture of bone marrow-derived macrophages from mice for functional analysis. STAR Protoc 2021;2:100246.
9. Schmittgen TD, Livak KJ. Analyzing real-time PCR data by the comparative C(T) method. Nat Protoc 2008;3:1101–8.
10. Harada Y, Miyamoto K, Sujino T. Protocol to isolate and enrich mouse splenic naive CD4(+) T cells for in vitro CD4(+)CD8alphaalpha(+) cell induction. STAR Protoc 2022;3:101728.
11. Hao Y, Stuart T, Kowalski MH, et al. Dictionary learning for integrative, multimodal and scalable single-cell analysis. Nat Biotechnol 2024;42:293–304.
12. Korsunsky I, Millard N, Fan J, et al. Fast, sensitive and accurate integration of single-cell data with Harmony. Nat Methods 2019;16:1289–1296.
13. Yu G, Wang LG, Han Y, et al. clusterProfiler: an R package for comparing biological themes among gene clusters. OMICS 2012;16:284–7.
14. Jin S, Guerrero-Juarez CF, Zhang L, et al. Inference and analysis of cell-cell communication using CellChat. Nature Communications 2021;12:1088.
15. Colaprico A, Silva TC, Olsen C, et al. TCGAbiolinks: an R/Bioconductor package for integrative analysis of TCGA data. Nucleic Acids Res 2016;44:e71.
16. Mounir M, Lucchetta M, Silva TC, et al. New functionalities in the TCGAbiolinks package for the study and integration of cancer data from GDC and GTEx. PLoS Comput Biol 2019;15:e1006701.
17. Windhager J, Zanotelli VRT, Schulz D, et al. An end-to-end workflow for multiplexed

image processing and analysis. Nat Protoc 2023;18:3565–3613.

18. Van Gassen S, Callebaut B, Van Helden MJ, et al. FlowSOM: Using self-organizing maps for visualization and interpretation of cytometry data. Cytometry A 2015;87:636–45.

19. Patrick E, Canete NP, Iyengar SS, et al. Spatial analysis for highly multiplexed imaging data to identify tissue microenvironments. Cytometry A 2023;103:593–599.

20. Steele NG, Carpenter ES, Kemp SB, et al. Multimodal Mapping of the Tumor and Peripheral Blood Immune Landscape in Human Pancreatic Cancer. Nat Cancer 2020;1:1097–1112.

#### Supplementary Table 1. TCGA metadata

| Sample_Ba | TAM_Signature_Score | TAM_CCP_Group | vital_status | days_to_death | days_to_last_follow_up | gender | tumor_grade_category | ajcc_pathologic_stage |
|---|---|---|---|---|---|---|---|---|
| TCGA-HV- | -1.199589585 | Low | Alive | NA | 978 | male | Three Tier | Stage IIB |
| TCGA-3A- | -1.242481346 | Low | Alive | NA | NA | male | NA | NA |
| TCGA-2L- | -1.328590869 | Low | Alive | NA | 1383 | male | Three Tier | Stage IIB |
| TCGA-3A- | -1.563492936 | Low | Alive | NA | 1103 | female | Four Tier | Stage IB |
| TCGA-3A- | -1.696846956 | Low | Alive | NA | 1942 | male | Four Tier | NA |
| TCGA-3A- | -1.717648913 | Low | Alive | NA | 1542 | female | Four Tier | Stage IB |
| TCGA-US- | -2.273947026 | Low | Alive | NA | 1216 | male | Four Tier | Stage IIA |
| TCGA-HZ- | -2.29102664 | Low | Dead | 661 | NA | male | NA | NA |
| TCGA-3A- | -2.312168867 | Low | Alive | NA | 998 | male | Four Tier | Stage IB |
| TCGA-FB- | -2.555622361 | Low | Dead | 485 | 485 | male | Not Reported | Stage IIB |
| TCGA-HZ- | -0.137040647 | Mid-Low | Dead | 518 | NA | male | NA | NA |
| TCGA-3A- | -0.154041485 | Mid-Low | Alive | NA | 1021 | female | Not Reported | Stage IA |
| TCGA-HZ- | -0.320572629 | Mid-Low | Dead | 151 | 151 | female | Not Reported | Stage IA |
| TCGA-RB- | -0.344603092 | Mid-Low | Alive | NA | 286 | male | Four Tier | Stage IIB |
| TCGA-F2- | -0.358736436 | Mid-Low | Alive | NA | 517 | male | Not Reported | Stage IIA |
| TCGA-3A- | -0.380464397 | Mid-Low | Alive | NA | 2084 | female | Four Tier | Stage IB |
| TCGA-2L- | -0.385540627 | Mid-Low | Dead | 103 | 103 | male | Three Tier | Stage IIB |
| TCGA-2L- | -0.474995439 | Mid-Low | Dead | 684 | 684 | male | Three Tier | Stage IIB |
| TCGA-2J- | -0.492506487 | Mid-Low | Dead | 277 | NA | male | NA | NA |
| TCGA-3A- | -0.496091086 | Mid-Low | Dead | 458 | NA | male | NA | NA |
| TCGA-HZ- | -0.515808842 | Mid-Low | Alive | NA | NA | male | NA | NA |
| TCGA-LB- | -0.518861722 | Mid-Low | Alive | NA | 379 | female | Four Tier | Stage IIA |
| TCGA-2J- | -0.526272401 | Mid-Low | Dead | 607 | 607 | male | Four Tier | Stage IIB |
| TCGA-IB- | -0.529573775 | Mid-Low | Dead | 41 | 41 | male | Four Tier | Stage IIB |
| TCGA-2J- | -0.530574612 | Mid-Low | Alive | NA | 440 | male | Four Tier | Stage IIB |
| TCGA-IB- | -0.574379824 | Mid-Low | Alive | NA | 666 | male | Four Tier | Stage IIB |
| TCGA-2J- | -0.590196461 | Mid-Low | Alive | NA | 484 | male | Four Tier | Stage IIB |
| TCGA-3E- | -0.596452816 | Mid-Low | Dead | 2182 | 2182 | male | Not Reported | Stage IIA |
| TCGA-2L- | -0.6236249 | Mid-Low | Dead | 143 | 143 | male | Three Tier | Stage IIB |
| TCGA-HV- | -0.656484107 | Mid-Low | Dead | 2036 | NA | female | NA | NA |
| TCGA-HZ- | -0.663210281 | Mid-Low | Alive | NA | 0 | male | Not Reported | Stage IIB |
| TCGA-US- | -0.70118151 | Mid-Low | Dead | 430 | 430 | male | Four Tier | Stage IIB |
| TCGA-HV- | -0.701568478 | Mid-Low | Dead | 128 | 128 | male | Three Tier | Stage IIA |
| TCGA-S4- | -0.744675123 | Mid-Low | Alive | NA | 737 | male | Three Tier | Stage IIB |
| TCGA-FB- | -0.772105767 | Mid-Low | Dead | 1130 | 1130 | male | Three Tier | Stage IIB |
| TCGA-LB- | -0.841414461 | Mid-Low | Dead | 393 | 393 | female | Not Reported | Stage IIB |
| TCGA-3A- | -0.851297263 | Mid-Low | Alive | NA | NA | female | NA | Stage 0a |
| TCGA-3A- | -0.862118115 | Mid-Low | Alive | NA | NA | male | NA | NA |
| TCGA-3A- | -0.926586617 | Mid-Low | Dead | 308 | 308 | male | Four Tier | Stage IIB |
| TCGA-HV- | -0.942874499 | Mid-Low | Dead | 532 | 532 | female | Three Tier | Stage IIB |
| TCGA-US- | -1.003638591 | Mid-Low | Dead | 12 | 12 | male | Four Tier | Stage IIB |
| TCGA-FB- | -1.011543921 | Mid-Low | Dead | 244 | NA | male | NA | NA |
| TCGA-FB- | -1.016598377 | Mid-Low | Dead | 153 | NA | female | NA | NA |
| TCGA-F2- | -1.03345903 | Mid-Low | Alive | NA | 295 | male | Four Tier | Stage IIB |
| TCGA-HZ- | -1.046512782 | Mid-Low | Alive | NA | 7 | male | Four Tier | Stage IB |
| TCGA-US- | -1.065601495 | Mid-Low | Dead | 511 | 511 | female | Four Tier | Stage IIB |
| TCGA-YY- | -1.068221277 | Mid-Low | Alive | NA | 2016 | female | Not Reported | Stage IIB |
| TCGA-IB- | -1.072616677 | Mid-Low | Dead | 250 | 250 | male | Not Reported | Stage IIB |

**Supplementary Table 2. PAAD TCGA RNA-seq TAM signature gene list**

| TAM signature gene expression | Low | Mid-Low | Mid-High | High |
|---|---|---|---|---|
| IRF8 | 2.603091 | 4.331811 | 4.700806 | 5.643474 |
| CCL2 | 2.040215 | 4.130447 | 4.800148 | 5.570849 |
| C1QC | 4.677733 | 5.427456 | 6.985428 | 7.758906 |
| GBP5 | -0.466935 | 1.650967 | 2.970422 | 4.154965 |
| HCST | -1.846059 | -1.246234 | -0.36323 | 0.41649 |
| LILRB4 | 0.911844 | 2.495661 | 3.870706 | 4.852138 |
| AIF1 | 2.621298 | 3.435113 | 4.655514 | 5.419417 |
| PSMB9 | 3.66761 | 5.10511 | 5.691315 | 5.857204 |
| GBP4 | 3.226164 | 4.553997 | 5.193495 | 5.7214 |
| GBP1 | 2.220695 | 4.491734 | 5.299744 | 5.677418 |
| HLA-DOA | 2.008236 | 3.968088 | 4.990232 | 6.095636 |
| C1QA | 4.978921 | 5.672427 | 7.141653 | 7.922386 |
| CCL4 | 0.530353 | 0.936317 | 1.972906 | 3.145736 |
| NCF1C | -2.014353 | -0.315412 | 0.4667 | 1.675827 |
| LAP3 | 5.450921 | 6.140024 | 6.431167 | 6.967938 |
| TNFAIP3 | 3.778375 | 5.316366 | 5.713401 | 6.404584 |
| ITGB2 | 3.734406 | 5.832964 | 6.932168 | 7.721272 |
| LAIR1 | 2.130115 | 3.492167 | 4.814516 | 5.739414 |
| FOLR2 | 2.001865 | 1.937361 | 3.556195 | 4.683393 |
| CD83 | 1.992903 | 2.944207 | 3.489717 | 4.564426 |
| SIGLEC1 | 0.86369 | 2.306506 | 3.962612 | 4.829061 |
| TCN2 | 4.573594 | 4.883537 | 5.247635 | 5.735209 |
| PLTP | 4.882175 | 6.028384 | 6.585692 | 7.070375 |
| C1QB | 4.952928 | 5.645017 | 7.239465 | 8.077531 |
| DOK2 | 1.245437 | 2.003021 | 3.107578 | 4.040649 |
| GIMAP6 | 2.674815 | 3.030215 | 3.763615 | 4.815818 |
| CD40 | 1.621133 | 4.601374 | 4.815802 | 5.225843 |
| CCL3 | -1.834573 | -1.78089 | -0.772755 | 0.065797 |
| CCL8 | -2.236897 | -1.863083 | -0.669797 | 0.533219 |
| FCN1 | -0.679927 | 0.585344 | 2.231447 | 3.966539 |
| CD4 | 3.577383 | 4.698165 | 5.776899 | 6.743999 |
| VAV1 | 1.099501 | 3.62298 | 3.947831 | 4.617501 |
| TLR7 | -0.989057 | 0.383815 | 1.741052 | 2.91165 |
| FGD2 | 1.003684 | 2.997072 | 3.650108 | 4.360607 |
| LST1 | 1.223471 | 2.485336 | 3.640827 | 4.462297 |
| VSIG4 | 1.983632 | 3.186976 | 4.86132 | 5.807427 |
| CLEC7A | 0.205349 | 2.055259 | 3.399553 | 4.191196 |

### Supplementary Table 3. Human PDA scRNA-seq cell count

| Patiet ID | AdjNorm_1 | | AdjNorm_2 | | AdjNorm_3 | | PDAC_1 | | PDAC_2 | | PDAC_3 | | PDAC_4 | |
|---|---|---|---|---|---|---|---|---|---|---|---|---|---|---|
| Celltype | count | proportion | count | proportion | count | proportion | count | proportion | count | proportion | count | proportion | count | proportion |
| Myeloid | 803 | 19.119048 | 819 | 28.368549 | 67 | 6.4237776 | 186 | 11.625 | 289 | 16.571101 | 205 | 9.255079 | 529 | 24.056389 |
| CD4 T cell | 66 | 1.5714286 | 5 | 0.1731902 | 1 | 0.0958773 | 90 | 5.625 | 30 | 1.7201835 | 496 | 22.392777 | 208 | 9.4588449 |
| CD8 T cell | 820 | 19.52381 | 509 | 17.630759 | 60 | 5.7526366 | 52 | 3.25 | 41 | 2.3509174 | 247 | 11.151242 | 218 | 9.9135971 |
| Granulocyte | 476 | 11.333333 | 49 | 1.6972636 | 1 | 0.0958773 | 267 | 16.6875 | 63 | 3.6123853 | 15 | 0.6772009 | 104 | 4.7294225 |
| Epithelial cell | 522 | 12.428571 | 61 | 2.11292 | 121 | 11.601151 | 481 | 30.0625 | 1080 | 61.926606 | 905 | 40.857788 | 463 | 21.055025 |
| Mast cell | 199 | 4.7380952 | 77 | 2.6671285 | 5 | 0.4793864 | 4 | 0.25 | 4 | 0.2293578 | 15 | 0.6772009 | 19 | 0.8640291 |
| Acinar | 413 | 9.8333333 | 1156 | 40.041566 | 700 | 67.114094 | 247 | 15.4375 | 5 | 0.2866972 | 0 | 0 | 203 | 9.2314688 |
| NK cell | 174 | 4.1428571 | 62 | 2.147558 | 7 | 0.6711409 | 237 | 14.8125 | 77 | 4.4151376 | 109 | 4.9209932 | 161 | 7.3215098 |
| Fibroblast | 43 | 1.0238095 | 3 | 0.1039141 | 23 | 2.2051774 | 1 | 0.0625 | 127 | 7.2821101 | 27 | 1.2189616 | 111 | 5.047749 |
| Plasma cell | 17 | 0.4047619 | 13 | 0.4502944 | 4 | 0.3835091 | 12 | 0.75 | 10 | 0.5733945 | 9 | 0.4063205 | 5 | 0.2273761 |
| B cell | 54 | 1.2857143 | 11 | 0.3810184 | 2 | 0.1917546 | 12 | 0.75 | 3 | 0.1720183 | 156 | 7.0428894 | 30 | 1.3642565 |
| Treg cell | 1 | 0.0238095 | 1 | 0.034638 | 0 | 0 | 7 | 0.4375 | 3 | 0.1720183 | 22 | 0.993228 | 2 | 0.0909504 |
| Pericyte | 462 | 11 | 29 | 1.0045029 | 1 | 0.0958773 | 4 | 0.25 | 5 | 0.2866972 | 3 | 0.1354402 | 40 | 1.8190086 |
| Endothelial cell | 150 | 3.5714286 | 92 | 3.186699 | 51 | 4.8897411 | 0 | 0 | 7 | 0.4013761 | 6 | 0.2708804 | 106 | 4.8203729 |
| total cells count | 4200 | | 2887 | | 1043 | | 1600 | | 1744 | | 2215 | | 2199 | |

| Patiet ID | PDAC_5 | | PDAC_6 | | PDAC_7 | | PDAC_8 | | PDAC_9 | | PDAC_10 | | PDAC_11 | |
|---|---|---|---|---|---|---|---|---|---|---|---|---|---|---|
| Celltype | count | proportion | count | proportion | count | proportion | count | proportion | count | proportion | count | proportion | count | proportion |
| Myeloid | 435 | 29.058116 | 129 | 6.1929909 | 1815 | 25.155925 | 699 | 15.897203 | 349 | 8.4238475 | 69 | 5.1918736 | 938 | 23.420724 |
| CD4 T cell | 59 | 3.9412158 | 236 | 11.329813 | 715 | 9.9099099 | 419 | 9.5292245 | 235 | 5.6722182 | 170 | 12.791573 | 46 | 1.1485643 |
| CD8 T cell | 52 | 3.4736139 | 160 | 7.681229 | 744 | 10.31185 | 720 | 16.374801 | 110 | 2.6550809 | 249 | 18.735892 | 51 | 1.2734082 |
| Granulocyte | 49 | 3.2732131 | 16 | 0.7681229 | 11 | 0.1524602 | 20 | 0.4548556 | 2570 | 62.032344 | 422 | 31.753198 | 8 | 0.1997503 |
| Epithelial cell | 496 | 33.132933 | 801 | 38.454153 | 181 | 2.5086625 | 877 | 19.945417 | 331 | 7.9893797 | 202 | 15.199398 | 2745 | 68.539326 |
| Mast cell | 8 | 0.5344021 | 39 | 1.8722996 | 1072 | 14.857935 | 329 | 7.4823743 | 20 | 0.482742 | 18 | 1.3544018 | 5 | 0.1248439 |
| Acinar | 131 | 8.750835 | 3 | 0.144023 | 0 | 0 | 8 | 0.1819422 | 132 | 3.186097 | 13 | 0.9781791 | 7 | 0.1747815 |
| NK cell | 122 | 8.1496326 | 206 | 9.8895823 | 121 | 1.6770617 | 188 | 4.2756425 | 302 | 7.2894038 | 38 | 2.8592927 | 16 | 0.3995006 |
| Fibroblast | 104 | 6.9472278 | 54 | 2.5924148 | 444 | 6.1538462 | 438 | 9.9613373 | 44 | 1.0620323 | 13 | 0.9781791 | 97 | 2.4219725 |
| Plasma cell | 7 | 0.4676019 | 14 | 0.6721075 | 888 | 12.307692 | 322 | 7.3231749 | 9 | 0.2172339 | 21 | 1.5801354 | 31 | 0.7740325 |
| B cell | 6 | 0.4008016 | 399 | 19.155065 | 82 | 1.1365211 | 122 | 2.7746191 | 24 | 0.5792904 | 37 | 2.7840482 | 19 | 0.474407 |
| Treg cell | 5 | 0.3340013 | 20 | 0.9601536 | 379 | 5.2529453 | 172 | 3.911758 | 7 | 0.1689597 | 69 | 5.1918736 | 11 | 0.2746567 |
| Pericyte | 3 | 0.2004008 | 2 | 0.0960154 | 610 | 8.4546085 | 46 | 1.0461678 | 2 | 0.0482742 | 4 | 0.3009782 | 3 | 0.0749064 |
| Endothelial cell | 20 | 1.3360053 | 4 | 0.1920307 | 153 | 2.1205821 | 37 | 0.8414828 | 8 | 0.1930968 | 4 | 0.3009782 | 28 | 0.6991261 |
| total cells count | 1497 | | 2083 | | 7215 | | 4397 | | 4143 | | 1329 | | 4005 | |

| Patiet ID | PDAC_11B | | PDAC_12 | | PDAC_13 | | PDAC_14 | | PDAC_15 | | PDAC_16 | |
|---|---|---|---|---|---|---|---|---|---|---|---|---|
| Celltype | count | proportion | count | proportion | count | proportion | count | proportion | count | proportion | count | proportion |
| Myeloid | 222 | 10.330386 | 947 | 59.33584 | 1193 | 24.008855 | 158 | 23.688156 | 964 | 33.765324 | 25 | 2.6399155 |
| CD4 T cell | 165 | 7.6779898 | 92 | 5.764411 | 301 | 6.0575569 | 292 | 43.778111 | 308 | 10.788091 | 24 | 2.5343189 |
| CD8 T cell | 32 | 1.4890647 | 53 | 3.320802 | 170 | 3.4212115 | 74 | 11.094453 | 174 | 6.0945709 | 9 | 0.9503696 |
| Granulocyte | 958 | 44.578874 | 55 | 3.4461153 | 320 | 6.4399276 | 36 | 5.3973013 | 142 | 4.9737303 | 0 | 0 |
| Epithelial cell | 616 | 28.664495 | 24 | 1.5037594 | 2733 | 55.001006 | 31 | 4.6476762 | 800 | 28.021016 | 827 | 87.328405 |
| Mast cell | 9 | 0.4187994 | 1 | 0.0626566 | 30 | 0.6037432 | 3 | 0.4497751 | 9 | 0.3152364 | 10 | 1.0559662 |
| Acinar | 4 | 0.1861331 | 314 | 19.674185 | 0 | 0 | 8 | 1.1994003 | 1 | 0.0350263 | 0 | 0 |
| NK cell | 58 | 2.6989297 | 75 | 4.6992481 | 138 | 2.7772188 | 28 | 4.197901 | 253 | 8.8616462 | 4 | 0.4223865 |
| Fibroblast | 17 | 0.7910656 | 0 | 0 | 47 | 0.9458644 | 5 | 0.7496252 | 120 | 4.2031524 | 32 | 3.3790919 |
| Plasma cell | 11 | 0.511866 | 11 | 0.6892231 | 3 | 0.0603743 | 4 | 0.5997001 | 28 | 0.9807356 | 5 | 0.5279831 |
| B cell | 29 | 1.3494649 | 12 | 0.7518797 | 13 | 0.2616221 | 21 | 3.1484258 | 18 | 0.6304729 | 4 | 0.4223865 |
| Treg cell | 13 | 0.6049325 | 9 | 0.5639098 | 17 | 0.3421212 | 4 | 0.5997001 | 31 | 1.0858144 | 4 | 0.4223865 |
| Pericyte | 11 | 0.511866 | 0 | 0 | 2 | 0.0402495 | 1 | 0.149925 | 2 | 0.0700525 | 2 | 0.2111932 |
| Endothelial cell | 4 | 0.1861331 | 3 | 0.1879699 | 2 | 0.0402495 | 2 | 0.2998501 | 5 | 0.1751313 | 1 | 0.1055966 |
| total cells count | 2149 | | 1596 | | 4969 | | 667 | | 2855 | | 947 | |

#### Supplementary Table 4. Mouse KPC tumor scRNA-seq cell count

| Total cell type | WT | | *Csf1r-Porcn-/-* | |
|---|---|---|---|---|
| Complete_Labels | Count | Proportion | Count | Proportion |
| Basal cancer cell | 948 | 0.079092274 | 535 | 0.047340943 |
| Classical cancer cell | 8688 | 0.724845653 | 5789 | 0.512255553 |
| CD8 T cell | 138 | 0.011513432 | 1102 | 0.097513494 |
| NK cell | 27 | 0.002252628 | 151 | 0.013361649 |
| CD4 T cell | 110 | 0.009177374 | 803 | 0.071055659 |
| CAF | 788 | 0.065743367 | 883 | 0.078134678 |
| DC | 448 | 0.03737694 | 503 | 0.044509335 |
| Mono/Mac | 328 | 0.027365259 | 514 | 0.045482701 |
| Neutrophil | 328 | 0.027365259 | 215 | 0.019024865 |
| B cell | 78 | 0.006507592 | 713 | 0.063091762 |
| Endothelial cell | 105 | 0.00876022 | 93 | 0.00822936 |
| | 11986 | | 11301 | |

| Mono/Mac subclustering | WT | | *Csf1r-Porcn-/-* | |
|---|---|---|---|---|
| Myeloid_labels | Count | Proportion_within | Count | Proportion_within_Myeloid |
| Trem2+ Macrophage | 137 | 0.417682927 | 121 | 0.246938776 |
| C1q+ Macrophage | 81 | 0.24695122 | 112 | 0.228571429 |
| Hexb+ Macrophage | 45 | 0.137195122 | 26 | 0.053061224 |
| Activated Monocyte | 50 | 0.152439024 | 153 | 0.312244898 |
| Classical Monocyte | 15 | 0.045731707 | 78 | 0.159183673 |

**Supplementary Table 5. Mouse scRNA-seq GO analysis**

| GOBP_Hexb mac GO(ONE VS REST) | | | | | | | | | |
|---|---|---|---|---|---|---|---|---|---|
| ID | Description | GeneRatio | BgRatio | pvalue | p.adjust | qvalue | geneID | Count | -log10(adj.p |
| GO:0060485 | mesenchyme development | 30/362 | 309/18539 | 5.06E-13 | 1.80E-09 | 1.33E-09 | Phldb2/Tcf7 | 30 | 8.7447275 |
| GO:0016055 | Wnt signaling pathway | 29/362 | 421/18539 | 4.60E-09 | 1.24E-06 | 9.11E-07 | Tcf7l2/Cdh2 | 29 | 5.9065783 |
| GO:0198738 | cell-cell signaling by wnt | 29/362 | 423/18539 | 5.11E-09 | 1.24E-06 | 9.13E-07 | Tcf7l2/Cdh2 | 29 | 5.9065783 |
| GO:0031018 | endocrine pancreas development | 10/362 | 45/18539 | 1.24E-08 | 2.43E-06 | 1.79E-06 | Cdh2/Foxa2 | 10 | 5.6143937 |
| GO:0009611 | response to wounding | 30/362 | 488/18539 | 3.30E-08 | 4.93E-06 | 3.63E-06 | Tmeff2/Par | 30 | 5.3071531 |
| GOBP_ C1q mac GO(ONE VS REST) | | | | | | | | | |
| GO:0050900 | leukocyte migration | 27/339 | 366/18539 | 8.90568366 | 8.81662682 | 6.86440722 | Cadm1/Cd8 | 27 | 6.0546975 |
| GO:0050853 | B cell receptor signaling pathway | 11/339 | 55/18539 | 3.79398375 | 2.44712234 | 1.90526883 | Prkcb/Cd81 | 11 | 5.6113443 |
| GO:0002274 | myeloid leukocyte activation | 20/339 | 248/18539 | 3.31196897 | 1.00887670 | 7.85486406 | C1qa/Tgfbr | 20 | 4.9961619 |
| GO:0046651 | lymphocyte proliferation | 22/339 | 335/18539 | 2.59468452 | 4.10998029 | 3.19992883 | Tgfbr2/Cd8 | 22 | 4.3861603 |
| GO:0050851 | antigen receptor-mediated signaling pathway | 15/339 | 174/18539 | 7.58681129 | 9.69153958 | 7.54559262 | Prkcb/Fyb/ | 15 | 4.0136072 |
| GOBP_Trem2 mac GO(ONE VS REST) | | | | | | | | | |
| GO:0061620 | glycolytic process through glucose-6-phosphate | 10/394 | 20/18539 | 2.56558040 | 9.98523894 | 7.57791434 | Pgk1/Aldoa | 10 | 9.0006415 |
| GO:0071456 | cellular response to hypoxia | 16/394 | 114/18539 | 2.47326779 | 2.75027379 | 2.08721487 | Pgk1/Adam | 16 | 6.5606241 |
| GO:0002548 | monocyte chemotaxis | 12/394 | 58/18539 | 2.65451421 | 2.86982481 | 2.17794353 | Lgals3/Ccl6 | 12 | 6.5421446 |
| GO:0048771 | tissue remodeling | 18/394 | 196/18539 | 2.14077205 | 1.28182843 | 9.72794558 | Spp1/Adam | 18 | 4.8921701 |
| GO:0042060 | wound healing | 22/394 | 350/18539 | 6.43013185 | 0.0002195 | 0.0001666 | Mmp12/Hm | 22 | 3.6585121 |

#### Supplementary Table 6. IMC double reporter mouse KPC tumor cell count matrix

| | CD45+ | | | | | | |
|---|---|---|---|---|---|---|---|
| | Cell type | Activated monocyte | Mono-derived mac 1 | Mono-derived mac 2 | Mono-derived mac 3 | Mono-derived mac 4 | Resident mac 1 | Resident mac 2 |
| Adj. T | Pancreas_001 | 496(7.43) | 463(6.94) | 299(4.48) | 1031(15.44) | 48(0.72) | 391(5.86) | 359(5.38) |
| | Pancreas_002 | 117(3.58) | 206(6.31) | 225(6.89) | 413(12.64) | 44(1.35) | 223(6.83) | 199(6.09) |
| Early T | Tumor_001 | 100(1.86) | 87(1.62) | 141(2.63) | 671(12.49) | 92(1.71) | 235(4.38) | 194(3.61) |
| | Tumor_002 | 85(1.60) | 106(2.00) | 72(1.36) | 245(4.62) | 88(1.66) | 346(6.52) | 187(3.52) |
| Late T | Tumor_003 | 8(0.17) | 11(0.23) | 13(0.28) | 68(1.45) | 861(18.38) | 103(2.20) | 186(3.97) |
| | Tumor_004 | 1(0.02) | 4(0.09) | 4(0.09) | 10(0.22) | 478(10.38) | 16(0.35) | 34(0.74) |

| | CD45+ | | | | | | | |
|---|---|---|---|---|---|---|---|---|
| | Cell type | Resident mac 3 | Resident mac 4 | Resident mac 5 | Resident mac 6 | Resident mac 7 | Neutrophil | B cell |
| Adj. T | Pancreas_001 | 187(2.80) | 43(0.64) | 417(6.25) | 202(3.03) | 68(1.02) | 34(0.51) | 11(0.16) |
| | Pancreas_002 | 78(2.39) | 19(0.58) | 270(8.26) | 89(2.72) | 36(1.10) | 6(0.18) | 4(0.12) |
| Early T | Tumor_001 | 643(11.97) | 107(1.99) | 184(3.43) | 544(10.13) | 74(1.38) | 392(7.30) | 20(0.37) |
| | Tumor_002 | 570(10.74) | 213(4.01) | 242(4.56) | 359(6.76) | 76(1.43) | 92(1.73) | 7(0.13) |
| Late T | Tumor_003 | 608(12.98) | 794(16.95) | 121(2.58) | 91(1.94) | 541(11.55) | 220(4.70) | 10(0.21) |
| | Tumor_004 | 94(2.04) | 1362(29.57) | 90(1.95) | 92(2.00) | 314(6.82) | 1885(40.92) | 39(0.85) |

| | CD45- | | | | | | | |
|---|---|---|---|---|---|---|---|---|
| | Cell type | Epithelial cell | Fibroblast 1 | Fibroblast 2 | Fibroblast 3 | SMA+ cell 1 | SMA+ cell 2 | Total cell count |
| Adj. T | Pancreas_001 | 360(5.39) | 541(8.10) | 507(7.59) | 713(10.68) | 408(6.11) | 98(1.47) | 6676 |
| | Pancreas_002 | 205(6.27) | 234(7.16) | 255(7.81) | 405(12.40) | 205(6.27) | 34(1.04) | 3267 |
| Early T | Tumor_001 | 191(3.56) | 264(4.92) | 362(6.74) | 492(9.16) | 516(9.61) | 62(1.15) | 5371 |
| | Tumor_002 | 257(4.84) | 640(12.06) | 927(17.46) | 376(7.08) | 229(4.31) | 191(3.60) | 5308 |
| Late T | Tumor_003 | 24(0.51) | 475(10.14) | 11(0.23) | 33(0.70) | 471(10.05) | 36(0.77) | 4685 |
| | Tumor_004 | 27(0.59) | 43(0.93) | 91(1.98) | 14(0.30) | 6(0.13) | 2(0.04) | 4606 |

**Supplementary Table 7. IMC *Csf1r-Porcn*$^{-/-}$ mouse KPC tumor cell count matrix**

| | Total cell type | Epi1 | | Epi2 | | Epi3 | | Epi4 | |
|---|---|---|---|---|---|---|---|---|---|
| | Cell type | Count | % of total c | Count | % of total c | Count | % of total c | Count | % of total c |
| WT | 250910_KPC_WT_4078_001 | 1867 | 26.292072 | 800 | 11.266019 | 469 | 6.6047036 | 539 | 7.5904802 |
| | 250910_KPC_WT_4078_002 | 2235 | 28.283979 | 557 | 7.0488484 | 405 | 5.1252847 | 911 | 11.528727 |
| | 250910_KPC_WT_4079_001 | 389 | 8.6502113 | 277 | 6.159662 | 167 | 3.7135868 | 504 | 11.207472 |
| | 250910_KPC_WT_4079_002 | 1184 | 19.432135 | 782 | 12.8344 | 408 | 6.6962088 | 1340 | 21.99245 |
| | 250910_KPC_WT_4079_003 | 815 | 14.626705 | 447 | 8.0222541 | 286 | 5.1328069 | 905 | 16.241924 |
| KO | 250910_KPC_KO_4082_001 | 1660 | 29.882988 | 431 | 7.7587759 | 486 | 8.7488749 | 910 | 16.381638 |
| | 250910_KPC_KO_4082_002 | 1790 | 29.377975 | 628 | 10.30691 | 528 | 8.6656819 | 796 | 13.064172 |
| | 250910_KPC_KO_4082_011 | 1354 | 22.369073 | 692 | 11.432348 | 435 | 7.1865191 | 698 | 11.531472 |
| | 250910_KPC_KO_4083_001 | 2655 | 32.17012 | 1253 | 15.182358 | 881 | 10.674906 | 662 | 8.0213256 |
| | 250910_KPC_KO_4083_002 | 2569 | 36.162725 | 1005 | 14.146959 | 605 | 8.5163288 | 387 | 5.4476351 |
| | 250910_KPC_KO_4083_004 | 1417 | 33.657957 | 684 | 16.247031 | 377 | 8.9548694 | 508 | 12.066508 |

| | Total cell type | Apoptotic cancer cell | | iCAF | | myCAF | | Endothelial cell | | CD4+ T cell | |
|---|---|---|---|---|---|---|---|---|---|---|---|
| | Cell type | Count | % of total c | Count | % of total c | Count | % of total c | Count | % of total c | Count | % of total c |
| WT | 250910_KPC_WT_4078_001 | 123 | 1.7321504 | 1433 | 20.180256 | 433 | 6.0977327 | 369 | 5.1964512 | 234 | 2.9612756 |
| | 250910_KPC_WT_4078_002 | 130 | 1.6451531 | 2134 | 27.005821 | 473 | 5.9858264 | 402 | 5.0873197 | 101 | 2.2459417 |
| | 250910_KPC_WT_4079_001 | 214 | 4.758728 | 1154 | 25.661552 | 547 | 12.163665 | 69 | 1.5343562 | 93 | 1.5263417 |
| | 250910_KPC_WT_4079_002 | 390 | 6.4007878 | 571 | 9.3714098 | 393 | 6.4500246 | 100 | 1.6412276 | 280 | 5.0251256 |
| | 250910_KPC_WT_4079_003 | 314 | 5.6353195 | 1233 | 22.1285 | 588 | 10.552764 | 119 | 2.1356784 | 119 | 2.1422142 |
| KO | 250910_KPC_KO_4082_001 | 224 | 4.0324032 | 376 | 6.7686769 | 510 | 9.1809181 | 46 | 0.8280828 | 178 | 2.9213852 |
| | 250910_KPC_KO_4082_002 | 319 | 5.2355162 | 404 | 6.6305597 | 294 | 4.8252093 | 61 | 1.0011489 | 404 | 6.6743763 |
| | 250910_KPC_KO_4082_011 | 331 | 5.4683628 | 881 | 14.554766 | 338 | 5.5840079 | 61 | 1.0077647 | 357 | 4.3256997 |
| | 250910_KPC_KO_4083_001 | 450 | 5.4525627 | 505 | 6.118987 | 254 | 3.0776687 | 39 | 0.4725554 | 227 | 3.1953829 |
| | 250910_KPC_KO_4083_002 | 290 | 4.0822072 | 427 | 6.0106982 | 331 | 4.6593468 | 41 | 0.5771396 | 172 | 4.0855107 |
| | 250910_KPC_KO_4083_004 | 279 | 6.6270784 | 249 | 5.9144893 | 130 | 3.087886 | 26 | 0.6175772 | 52 | 1.2351544 |

| | Total cell type | CTL | | Proliferating CD4+ T cell | | Proliferating CD8+ T cell | | B cell | |
|---|---|---|---|---|---|---|---|---|---|
| | Cell type | Count | % of total c | Count | % of total cells | Count | % of total cells | Count | % of total cells |
| WT | 250910_KPC_WT_4078_001 | 229 | 3.2248979 | 56 | 0.788621321 | 31 | 0.436558231 | 3 | 0.042247571 |
| | 250910_KPC_WT_4078_002 | 245 | 3.1004809 | 7 | 0.088585168 | 24 | 0.303720577 | 1 | 0.012655024 |
| | 250910_KPC_WT_4079_001 | 108 | 2.4016011 | 16 | 0.355792751 | 5 | 0.111185235 | 1 | 0.022237047 |
| | 250910_KPC_WT_4079_002 | 264 | 4.332841 | 28 | 0.459543739 | 27 | 0.443131462 | 14 | 0.229771869 |
| | 250910_KPC_WT_4079_003 | 160 | 2.8715004 | 6 | 0.107681263 | 17 | 0.305096913 | 5 | 0.089734386 |
| KO | 250910_KPC_KO_4082_001 | 178 | 3.2043204 | 17 | 0.306030603 | 36 | 0.648064806 | 84 | 1.512151215 |
| | 250910_KPC_KO_4082_002 | 320 | 5.2519284 | 28 | 0.459543739 | 47 | 0.77137699 | 41 | 0.672903332 |
| | 250910_KPC_KO_4082_011 | 423 | 6.9882703 | 22 | 0.363456137 | 75 | 1.239055014 | 46 | 0.759953742 |
| | 250910_KPC_KO_4083_001 | 565 | 6.8459954 | 15 | 0.18175209 | 86 | 1.042045317 | 0 | 0 |
| | 250910_KPC_KO_4083_002 | 491 | 6.9115991 | 16 | 0.225225225 | 83 | 1.168355856 | 17 | 0.239301802 |
| | 250910_KPC_KO_4083_004 | 200 | 4.7505938 | 3 | 0.071258907 | 39 | 0.926365796 | 0 | 0 |

| | Macrophage subtype | CCR2+mono | | IL1b+mono | | CD206+mac | | Hexb+mac1 | |
|---|---|---|---|---|---|---|---|---|---|
| | Cell type | Count | % of total c | Count | % of total c | Count | % of total c | Count | % of total c |
| WT | 250910_KPC_WT_4078_001 | 29 | 0.4083932 | 4 | 0.0563301 | 44 | 0.619631 | 122 | 1.7180679 |
| | 250910_KPC_WT_4078_002 | 67 | 0.8478866 | 5 | 0.0632751 | 14 | 0.1771703 | 15 | 0.1898254 |
| | 250910_KPC_WT_4079_001 | 30 | 0.6671114 | 23 | 0.5114521 | 21 | 0.466978 | 366 | 8.1387592 |
| | 250910_KPC_WT_4079_002 | 9 | 0.1477105 | 48 | 0.7877893 | 3 | 0.0492368 | 26 | 0.4267192 |
| | 250910_KPC_WT_4079_003 | 8 | 0.143575 | 24 | 0.4307251 | 9 | 0.1615219 | 29 | 0.5204594 |
| KO | 250910_KPC_KO_4082_001 | 0 | 0 | 22 | 0.3960396 | 1 | 0.0180018 | 31 | 0.5580558 |
| | 250910_KPC_KO_4082_002 | 19 | 0.3118333 | 56 | 0.9190875 | 2 | 0.0328246 | 57 | 0.9354998 |
| | 250910_KPC_KO_4082_011 | 14 | 0.2312903 | 56 | 0.9251611 | 29 | 0.4791013 | 7 | 0.1156451 |
| | 250910_KPC_KO_4083_001 | 9 | 0.1090513 | 97 | 1.1753302 | 3 | 0.0363504 | 28 | 0.3392706 |
| | 250910_KPC_KO_4083_002 | 6 | 0.0844595 | 168 | 2.3648649 | 0 | 0 | 74 | 1.0416667 |
| | 250910_KPC_KO_4083_004 | 4 | 0.0950119 | 83 | 1.9714964 | 1 | 0.023753 | 23 | 0.5463183 |

| | Macrophage subtype | Hexb+mac2 | | Trem2+mac1 | | Trem2+mac2 | | Arg1+mac | | Neutrophil | | |
|---|---|---|---|---|---|---|---|---|---|---|---|---|
| | Cell type | Count | % of total c | Count | % of total c | Count | % of total c | Count | % of total c | Count | % of total c | Total cell cc |
| WT | 250910_KPC_WT_4078_001 | 273 | 3.8445289 | 5 | 0.0704126 | 0 | 0 | 7 | 0.0985777 | 31 | 0.4365582 | 7101 |
| | 250910_KPC_WT_4078_002 | 138 | 1.7463933 | 0 | 0 | 0 | 0 | 29 | 0.3669957 | 9 | 0.1138952 | 7902 |
| | 250910_KPC_WT_4079_001 | 107 | 2.379364 | 91 | 2.0235713 | 123 | 2.7351568 | 63 | 1.400934 | 129 | 2.8685791 | 4497 |
| | 250910_KPC_WT_4079_002 | 130 | 2.1335959 | 22 | 0.3610701 | 7 | 0.1148859 | 20 | 0.3282455 | 47 | 0.771377 | 6093 |
| | 250910_KPC_WT_4079_003 | 98 | 1.758794 | 83 | 1.4895908 | 4 | 0.0717875 | 211 | 3.7867911 | 92 | 1.6511127 | 5572 |
| KO | 250910_KPC_KO_4082_001 | 112 | 2.0162016 | 191 | 3.4383438 | 3 | 0.0540054 | 24 | 0.4320432 | 35 | 0.630063 | 5555 |
| | 250910_KPC_KO_4082_002 | 96 | 1.5755785 | 103 | 1.6904645 | 52 | 0.8534384 | 21 | 0.3446578 | 27 | 0.4431315 | 6093 |
| | 250910_KPC_KO_4082_011 | 34 | 0.5617049 | 131 | 2.1642161 | 10 | 0.1652073 | 17 | 0.2808525 | 42 | 0.6938708 | 6053 |
| | 250910_KPC_KO_4083_001 | 74 | 0.8966436 | 356 | 4.3135829 | 4 | 0.0484672 | 20 | 0.2423361 | 70 | 0.8481764 | 8253 |
| | 250910_KPC_KO_4083_002 | 79 | 1.1120495 | 222 | 3.125 | 8 | 0.1126126 | 38 | 0.5349099 | 75 | 1.0557432 | 7104 |
| | 250910_KPC_KO_4083_004 | 63 | 1.4964371 | 14 | 0.3325416 | 1 | 0.023753 | 0 | 0 | 57 | 1.3539192 | 4210 |

**Supplementary Table 8. PCR Oligonucleotide Sequences**

| qPCR Oligonucleotide Sequences | | | |
|---|---|---|---|
| Target | Forward | Reverse | Citation/Source |
| Wnt1 | ATCCATCTCTCCCACCTCCTAC | GAATCTTTCTCTCACCCTCTGG | |
| Wnt2 | GTGATGTGTGACAATGTGCCA | GTTGCAGTTCCAGCGATGC | |
| Wnt3 | AGCGTAGCAGAAGGTGTGAAG | CCAGGTGGCCCCTTATGATG | |
| Wnt4 | AGAACTGGAGAAGTGTGGCTGT | AAAGGACTGTGAGAAGGCTACG | |
| Wnt5a | GTCCTTTGAGATGGGTGGTATC | ACCTCTGGGTTAGGGAGTGTCT | |
| Wnt5b | CCCTGTGGAAAGACCTGATAAC | CTTGACAGAGTGACAGCAGATT | |
| Wnt6 | TTTACACCAGCCCACGAAAG | ACTCACCCATCCATCCCAGTA | |
| Wnt7b | TGAAGCTGGAATGTAAGTGTCAC | CGCTGCGTTGTACTTCTCCT | |
| Wnt8a | ACGGTGGAATTGTCCTGAGCATG | GATGGCAGCAGAGCGGATGG | |
| Wnt9a | ATGGTGTGTCTGGCTCCTG | CAGTGGCTTCATTGGTAGTGCT | |
| Wnt10a | TCCTGTTCTTCCTACTGCTGCT | ACGCACACACACCTCCATC | |
| Wnt11 | CCCTGGAAACGAAGTGTAAATG | AGGTAGCGGGTCTTGAGGTC | |
| Porcn | GCTGTCTCCTGCCTACTGTCCA | TGCTTGCATGCTTCAGGTAAGA | Saha et al., 2016 |
| Arg1 | CTGGAACCCAGAGAGAGCAT | CTCCTCGAGGCTGTCCTTT | |

| Genotyping Oligonucleotide Sequences | | | |
|---|---|---|---|
| Target | Forward | Reverse | Citation/Source |
| Porcn | CAAGTCCTTCCGCATATGGT | TGCATGCTTCAGGTAAGACG | IMSR_JAX:020994 |
| Csf1r_Transgene | GTGAGCCAGGGGACATCG | ACACCGGCCTTATTCCAAG | IMSR_JAX:029206 |
| Csf1r_Internal Positive | CTAGGCCACAGAATTGAAAGATCT | GTAGGTGGAAATTCTAGCATCATCC | |
| Ms4a3_Common | AGAGAAATCATCAGGGCAGAAAT | | IMSR_JAX:036382 |
| Ms4a3_Wild Type | | GAAAGGGGAACAAGCGAAGAT | |
| Ms4a3_Mutant | | TTGGCGAGAGGGGAAAGAC | |
| Ai14_Wild Type | AAGGGAGCTGCAGTGGAGTA | CCGAAAATCTGTGGGAAGTC | IMSR_JAX:007914 |
| Ai14_Mutant | CTGTTCCTGTACGGCATGG | GGCATTAAAGCAGCGTATCC | |
| hCD68_Transgene | CGGCTCTGTGAATGACAATG | TTGGCAGTTGTGGCAAGTAG | IMSR_JAX:026827 |
| hCD68_Internal Positive | CTAGGCCACAGAATTGAAAGATCT | GTAGGTGGAAATTCTAGCATCATCC | |
| Csf1r-icre_Transgene | AGATGCCAGGACATCAGGAACCTG | ATCAGCCACACCAGACACAGAGATC | IMSR_JAX:019098 |
| Csf1r-icre_Internal Positive | CTAGGCCACAGAATTGAAAGATCT | GTAGGTGGAAATTCTAGCATCATCC | |

**Supplementary Table 9. Materials**

| Flow Cytometry Antibody | Clone | Source | Catalog Number |
|---|---|---|---|
| Purified CD16/32 | 93 | Biolegend | 101302 |
| CD45 BV650 | 30-F11 | Biolegend | 103151 |
| CD45 BUV395 | 30-F11 | BD Horizon | 564279 |
| CD45 FITC | 30-F11 | eBioscience | 11-0451-85 |
| CD11b APC | M1-70 | eBioscience | 17-0112-82 |
| CD11b eFluor450 | M1-70 | eBioscience | 48-0112-82 |
| CD11b APC-R700 | M1-70 | BD Biosciences | 564985 |
| CD64 PerCP-eFluor710 | X54-5/7.1 | eBioscience | 46-0641-82 |
| CD206 BV605 | C068C2 | Biolegend | 141721 |
| CD11c BV510 | N418 | Biolegend | 117337 |
| CD3 FITC | 17A2 | Biolegend | 100203 |
| CD3 PE-Cy7 | 17A2 | Biolegend | 100220 |
| CD3 BV395 | 17A2 | BD Biosciences | 569614 |
| CD8 APC-Cy7 | 53-6.7 | Biolegend | 100714 |
| CD4 PE | GK1.5 | eBioscience | 12-0041-82 |
| CD4 BV785 | GK1.5 | Biolegend | 100453 |
| CD25 PE-Cy5 | PC61 | Biolegend | 102010 |
| CD206 AF700 | C068C2 | Biolegend | 141734 |
| F4/80 PE-Cy7 | BM8 | Biolegend | 123114 |
| F4/80 PE | BM8 | eBioscience | 12-4801-82 |
| MHCII BV785 | M5/114.15.2 | Biolegend | 107645 |
| MHCII APC | M5/114.15.2 | eBioscience | 17-5321-82 |
| Ly6C PE-Cy7 | HK1.4 | Biolegend | 128018 |
| Ly6C FITC | HK1.4 | Biolegend | 128006 |
| Ly6G BV711 | 1A8 | Biolegend | 127643 |
| CD45R/B220 PE-Cy5 | RA3-6B2 | BD Pharmingen | 553091 |
| GranzymeB PE-Cy7 | QA18A28 | Biolegend | 396410 |
| FOXP3 eFluor450 | FJK-16s | eBioscience | 48-5773-82 |
| Arginase1 APC | A1exF5 | eBioscience | 17-3697-82 |
| TNF-α BV510 | MP6-XT22 | Biolegend | 506339 |
| IL-10 BV605 | JES5-16E3 | Biolegend | 505031 |
| IL-10 FITC | JES5-16E3 | eBioscience | 11-7101-41 |

| Live/Dead | Source | Catalog Number |
|---|---|---|
| DAPI | ThermoFisher Scientific | 62248 |

| Immunofluorescence/Immunohistochemistry | Clone | Source | Catalog Number |
|---|---|---|---|
| Anti-CD8α | EPR21769 | Abcam | ab217344 |
| Anti-F4/80 | CI:A3-1 | Abcam | ab6640 |
| CD11c | N418 | Invitrogen | 14-0114-82 |
| Anti-CCR2 | EPR20844-15 | Abcam | ab273050 |
| Anti-β-Catenin | 14/Beta-Catenin | BD Transduction Laboratories | 610153 |
| Non-phospho β-catenin (Ser33/37/Thr41) | D13A1 | Cell signaling | 8814S |
| Anti-p53 | Polyclonal | Abcam | ab131442 |
| Anti-Ki67 | D3B5 | Cell signaling | 12202S |
| Anti-Pan cytokeratin | AE1/AE3 + 5D3 | Abcam | ab86734 |
| Alexa Fluor 488 anti-Rat IgG (H+L) | Polyclonal | Invitrogen | A-11006 |
| Alexa Fluor 555 anti-Rat IgG (H+L) | Polyclonal | Invitrogen | A-21434 |
| Alexa Fluor 647 anti-Rat IgG (H+L) | Polyclonal | Invitrogen | A-21247 |
| Alexa Fluor 488 anti-Rabbit IgG (H+L) | Polyclonal | Invitrogen | A-11034 |
| Alexa Fluor 555 anti-Rabbit IgG (H+L) | Polyclonal | Invitrogen | A-31572 |
| Alexa Fluor 647 anti-Rabbit IgG (H+L) | Polyclonal | Invitrogen | A-31573 |
| Alexa Fluor 488 anti-Mouse IgG (H+L) | Polyclonal | Invitrogen | A-11001 |
| Alexa Fluor 555 anti-Mouse IgG (H+L) | Polyclonal | Invitrogen | A-31570 |
| Alexa Fluor 647 anti-Mouse IgG (H+L) | Polyclonal | Invitrogen | A-31571 |
| Alexa Fluor 488 anti-Armenian Hamster IgG (H+L) | Polyclonal | Invitrogen | A-78963 |
| Anti-Rabbit, HRP | | Dako | K400311-2 |
| Anti-Mouse, HRP | | Dako | K400111-2 |
| DAB | | Dako | K346711-2 |

| In situ hybridization | Clone | Source | Catalog Number |
|---|---|---|---|
| Anti-Human CD68 | KP1 | Abcam | ab955 |
| Anti-Human CD8 | CAL66 | Abcam | ab251596 |
| Anti-Human Hs-WNT5A | | ACD Bio | 604921 |

| In vivo CD8 T cell depletion | Clone | Source | Catalog Number |
|---|---|---|---|
| Anti-Mouse CD8α | 2.43 | Bioxcell | BE0061 |
| Rat IgG2b isotype control | LTF-2 | Bioxcell | BE0090 |

| Western Blot Antibody | Clone | Source | Catalog Number |
|---|---|---|---|
| Phospho-c-Jun (Ser63) | 54B3 | Cell signaling | 2361S |
| c-Jun | 60A8 | Cell signaling | 9165S |
| p-SAPK/JNK (Thr183/Tyr185) | Polyclonal | Cell signaling | 9251S |
| SAPK/JNK | Polyclonal | Cell signaling | 9252S |
| Non-phospho β-catenin (Ser33/37/Thr41) | D13A1 | Cell signaling | 8814S |
| Anti-β-Catenin | 14/Beta-Catenin | BD Transduction Laboratories | 610153 |
| Arginase-1 | D4E3M | Cell signaling | 93668S |
| Beta Actin | C4 | Santa Cruz | sc-47778 |
| Anti-mouse IgG, HRP | | Cell signaling | 7076P2 |
| Anti-Rabbit IgG, HRP | | Invitrogen | 31460 |

**Supplementary Table 10. IMC antibodies and concentrations for double reporter KPC tumor (fixed frozen)**

| Metal Tag | Targets | Clone | Catalog Number | Vendor | Dilution |
|---|---|---|---|---|---|
| 141Pr | aSMA | 1A4 | 3141017D | Standard BioTools | 1:500 |
| 142Nd | CD11c | N418 | 3209005B | Standard BioTools | 1:500 |
| 143Nd | MPO | EPR20257 | ab221847 | Abcam | 1:500 |
| 144Nd | | | | | |
| 145Nd | CD4 | RM4-5 | 3145002B | Standard BioTools | 1:500 |
| 146Nd | F4/80 | BM8 | 3146008B | Standard BioTools | 1:500 |
| 147Sm | CD45.2 | 1900-04-13 | 3147004B | Standard BioTools | 1:500 |
| 148Nd | CD68 | E3O7V | 29176SF | Cell signaling | 1:500 |
| 149Sm | | | | | |
| 150Nd | CD11c | AP-MAB0806 | NB110-97871 | Novus | 1:500 |
| 151Eu | CD64 | X54-5/7.1 | 3151012B | Standard BioTools | 1:500 |
| 152Sm | CD3e | 145-2C11 | 3152004B | Standard BioTools | 1:500 |
| 153Eu | PanCK | C-11 | ab264485 | Abcam | 1:500 |
| 154Sm | CD11b | M1/70 | 3154006B | Standard BioTools | 1:500 |
| 155Gd | CD25 | 3C7 | 101902 | Biolegend | 1:500 |
| 156Gd | Arg1 | Polyclonal | NBP1-32731 | Novus | 1:500 |
| 158Gd | MHCII | M5/114.15.2 | 107637 | Biolegend | 1:500 |
| 159Tb | | | | | |
| 160Gd | CD45R(B220) | RA3-6B2 | 3160012B | Standard BioTools | 1:500 |
| 161Dy | MHCII | M5/114.15.2 | NBP1-43312 | Novus | 1:500 |
| 162Dy | Ly6C | HK1.4 | 3162014B | Standard BioTools | 1:250 |
| 163Dy | CD8 | 53-6.7 | 100702 | Biolegend | 1:500 |
| 164Dy | TREM2 | EPR26210-1 | ab305072 | Abcam | 1:250 |
| 165Ho | CD31 | 1901-01-24 | 3165013B | Standard BioTools | 1:500 |
| 166Er | EpCAM | EPR20532-222 | ab232539 | Abcam | 1:500 |
| 167Er | Nkp46 | 29A1.4 | 3167008B | Standard BioTools | 1:500 |
| 168Er | Ki67 | B56 | 3168022D | Standard BioTools | 1:500 |
| 169Tm | CD206 | C068C2 | 3165013B | Standard BioTools | 1:500 |
| 170Er | | | | | |
| 171Yb | CCR2 | EPR20844-15 | ab273061 | abcam | 1:500 |
| 172Yb | CD86 | GL1 | 3172016B | Standard BioTools | 1:500 |
| 173Yb | HEXB | E9X5S | 28112SF | Cell signaling | 1:250 |
| 174Yb | CX3CR1 | Polyclonal | NBP1-76949 | Novus | 1:500 |
| 175Lu | MRC1(CD206) | EPR25215-277 | ab300622 | Abcam | 1:500 |
| 176Yb | PDGFRa | EPR22059-270 | ab234965 | Abcam | 1:500 |
| 191Ir | DNA1 | | | Standard BioTools | 1:400 |
| 193Ir | DNA2 | | | Standard BioTools | 1:400 |

**Supplementary Table 11. IMC antibodies and concentrations for *Csf1r-Porcn*$^{-/-}$ KPC tumor (FFPE)**

| Metal Tag | Targets | Clone | Catalogue Number | Vendor | Dilution |
|---|---|---|---|---|---|
| 141Pr | aSMA | 1A4 | 3141017D | Standard BioTools | 1:1600 |
| 142Nd | CD45 | 30-F11 | 14-0451-82 | Invitrogen | 1:200 |
| 143Nd | Vimentin | D21H3 | 3143027D | Standard BioTools | 1:1600 |
| 144Nd | beta-catenin | E247 | ab196204 | Abcam | 1:800 |
| 145Nd | CD31 | EPR17259 | ab225883 | Abcam | 1:800 |
| 146Nd | CD11c | D1V9Y | CST; 39143SF | Cell signaling | 1:200 |
| 147Sm | | | | | |
| 148Nd | CD68 | E3O7V | 29176SF | Cell signaling | 1:800 |
| 149Sm | CD11b | EPR1344 | 3149028D | Standard BioTools | 1:400 |
| 150Nd | non-p-b-cat(S37/T41) | EPR23969-131 | ab278503 | Abcam | 1:200 |
| 151Eu | phospho-c-Jun(Ser63) | Y172 | ab227533 | Abcam | 1:200 |
| 152Sm | WNT5A | Polyclonal | NBP2-24752 | Novus | 1:200 |
| 153Eu | Pan-CK | C-11 | ab264485 | Abcam | 1:200 |
| 154Sm | p-JNK(T183+Y185) | 81E11 | 58328 | Cell signaling | 1:200 |
| 155Gd | GranzymeB | EPR22645-206 | ab255868 | Abcam | 1:400 |
| 156Gd | Arginase 1 | Polyclonal | NBP1-32731 | Novus | 1:800 |
| 158Gd | MPO | EPR17996 | ab236022 | Abcam | 1:800 |
| 159Tb | CD4 | 4SM95 | 14-9766-82 | Invitrogen | 1:800 |
| 160Gd | CD45R(B220) | RA3-6B2 | 3160012B | Standard BioTools | 1:800 |
| 161Dy | MHCII | M5/114.15.2 | 14-5321-82 | Invitrogen | 1:400 |
| 162Dy | CD8 | EPR21769 | ab230156 | Abcam | 1:800 |
| 163Dy | IL-1β | 3A6 | 27989SF | Cell signaling | 1:400 |
| 164Dy | Trem2 | EPR26210-1 | ab305072 | Abcam | 1:200 |
| 165Ho | Nos2/iNOS | Polyclonal | NB300-605 | Novus | 1:400 |
| 166Er | EpCAM | EPR20532-222 | ab232539 | Abcam | 1:800 |
| 167Er | VEGFA | VG1 | NB100-664 | Novus | 1:400 |
| 168Er | Ki67 | B56 | 3168022D | Standard BioTools | 1:800 |
| 169Tm | COL1A1 | 3G3 | ab88147 | Abcam | 1:800 |
| 170Er | CD3e | CD3-12 | ab255972 | Abcam | 1:800 |
| 171Yb | CCR2 | EPR20844-15 | ab273061 | Abcam | 1:800 |
| 172Yb | Cleaved Caspase 3 | 5A1E | 94530SF | Cell signaling | 1:400 |
| 173Yb | HEXB | E9X5S | 28112SF | Cell signaling | 1:200 |
| 174Yb | CX3CR1 | Polyclonal | NBP1-76949 | Novus | 1:200 |
| 175Lu | MRC1(CD206) | EPR25215-277 | ab300622 | Abcam | 1:800 |
| 176Yb | PDGFRa | EPR22059-270 | ab234965 | Abcam | 1:400 |
| 191Ir | DNA1 | | | Standard BioTools | 1:400 |
| 193Ir | DNA2 | | | Standard BioTools | 1:400 |

**Supplementary Figures**

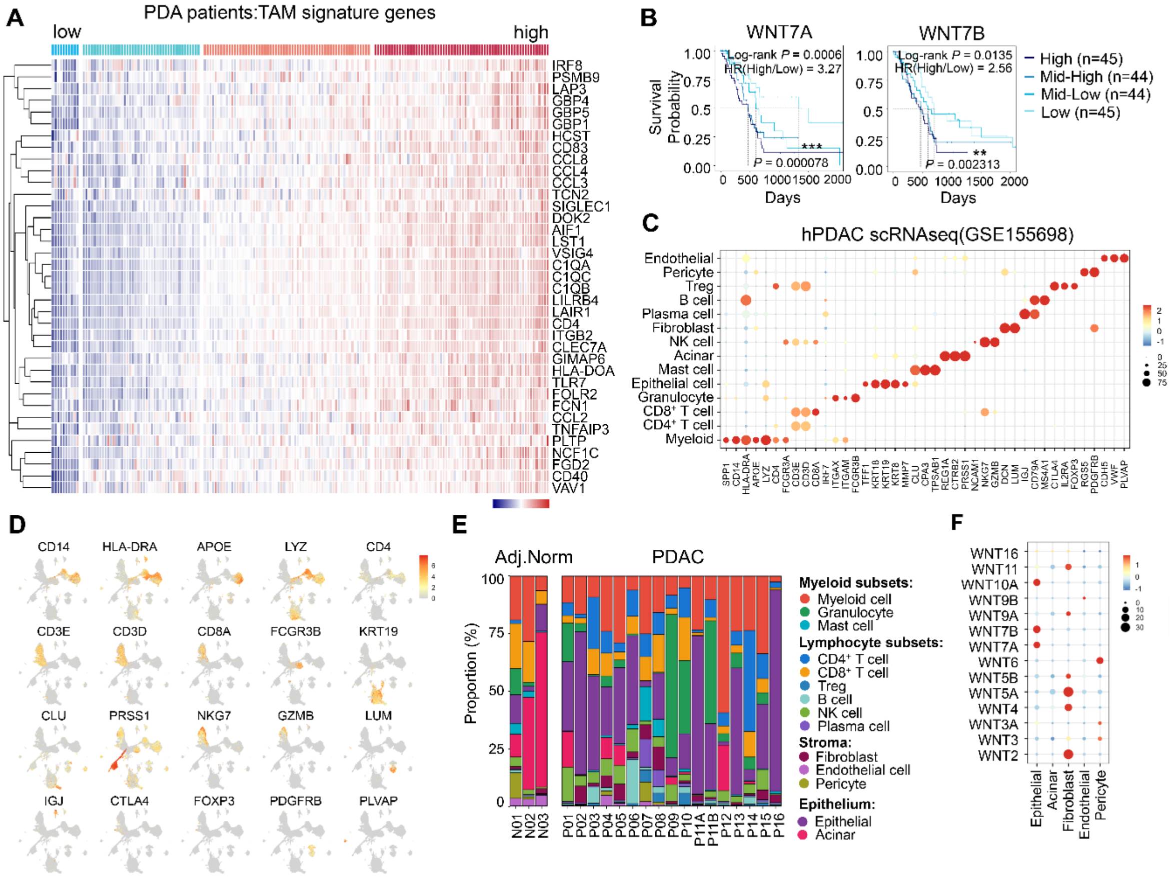

**Supplementary Figure S1. Transcriptomic evidence linking TAM enrichment to noncanonical WNT signaling in human PDAC.** (**A**) Heatmap of scaled expression of the 37-gene TAM signature across TCGA PDAC patients (n=178), hierarchically clustered by signature genes (rows) and patients (columns). (**B**) Kaplan–Meier overall survival curves for TCGA PDAC patients stratified into quartiles based on WNT7A (left) or WNT7B (right) expression (Low, Mid-low, Mid-high, High; n=45, 44, 44, 45, respectively). P values were calculated using the log-rank test. (**C**) Dot plot of marker-gene expression across major cell types from a published human PDAC scRNA-seq cohort (GSE155698; 17 PDAC tumors and three normal pancreas samples). Dot size indicates the fraction of cells expressing each gene, and color denotes scaled average expression. (**D**) UMAP feature plots showing expression of representative lineage markers used for cell-type annotation in the integrated scRNA-seq dataset. (**E**) Stacked bar plots showing per-sample cellular composition in adjacent normal pancreas (Adj. Norm; n=3) and PDAC tumors (n=17), based on the annotated scRNA-seq cell types. **(F)** Dot plot summarizing expression of WNT ligands across epithelial and stromal compartments in the scRNA-seq dataset. Dot size indicates the percentage of expressing cells; color indicates scaled average expression.

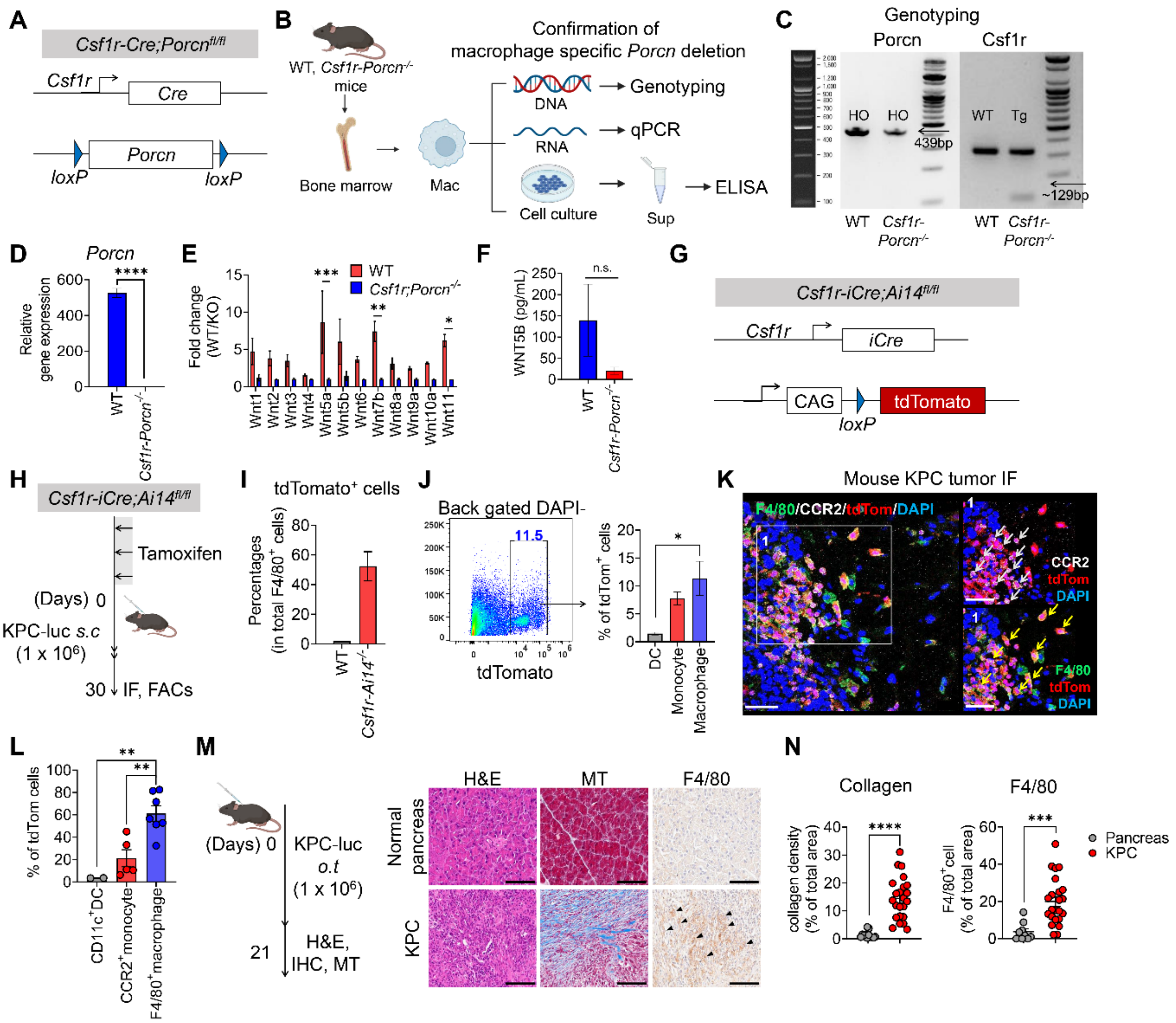

**Supplementary Figure S2. Characterization of *Csf1r-Porcn*$^{-/-}$ mice.** (**A**) Schematic of the conditional Porcn allele and Csf1r-Cre mating strategy used to ablate PORCN in Csf1r$^{+}$ myeloid cells (*Csf1r-Porcn*$^{-/-}$). (**B**) Experimental workflow to validate macrophage-specific Porcn deletion and loss of secreted WNTs. BMDMs were generated from WT and *Csf1r-Porcn*$^{-/-}$ mice and assessed by genomic PCR, qPCR, and ELISA. (**C**) Representative genotyping PCR confirming the presence of the Porcn floxed allele and Csf1r-Cre transgene in the indicated mice. (**D**) qPCR quantification of Porcn mRNA in WT versus *Csf1r-Porcn*$^{-/-}$ BMDMs. (**E**) qPCR profiling of Wnt ligand transcripts in WT and *Csf1r-Porcn*$^{-/-}$ BMDMs, shown as fold change (WT/KO). (**F**) ELISA measurement of WNT secretion in BMDM culture supernatants (shown as WNT5B concentration, pg/mL) from WT and *Csf1r-Porcn*$^{-/-}$ macrophages. (**G**)

Schematic of the tamoxifen-inducible lineage-tracing system used to assess Csf1r$^{+}$ cell targeting in tumors (Csf1r-iCre-Ai14tdt). (**H**) Timeline of orthotopic implantation of KPC cells and tamoxifen administration in Csf1r-iCre-Ai14$^{tdt}$ mice, followed by tumor harvest for IF and flow cytometry. (**I**) Quantification of tdTomato$^{+}$ cells among total F4/80$^{+}$ cells in tumors from WT versus Csf1r-iCre-Ai14$^{tdt}$ mice. (**J**) Representative flow-cytometry plot showing tdTomato labeling in tumor single-cell suspensions (back-gated on DAPI$^{-}$ live cells). (**K**) Representative tumor IF images showing tdTomato$^{+}$Csf1r-lineage cells co-localizing with myeloid markers in orthotopic KPC tumors. White arrows indicate CCR2$^{+}$tdTomato$^{+}$ monocytes and yellow arrows indicate F4/80$^{+}$tdTomato$^{+}$macrophages. Scale bars, 50 µm. (**L**) Quantification of tdTomato labeling across indicated immune populations within tumors (e.g., CD11c$^{+}$ DCs, CCR2$^{+}$ monocytes, and F4/80$^{+}$ macrophages). (**M**) Timeline of orthotopic KPC tumor harvest for histology (left). Representative H&E, Masson's trichrome (MT), and F4/80 IHC comparing normal pancreas and KPC tumors (right). Black arrows indicate F4/80$^{+}$macrophages. Scale bars, 100 µm. (**N**) Quantification of collagen density (MT-positive area) and F4/80$^{+}$ macrophage area in normal pancreas versus KPC tumors. The data are shown as means +/± S.E.M. For (**E**), Two-way ANOVA test, For (J,L), One-way ANOVA, For (N), Two-tailed unpaired t-test, * $p$ <0.05, ** $p$ <0.01, *** $p$ <0.001, **** $p$ <0.0001. n.s., not significant.

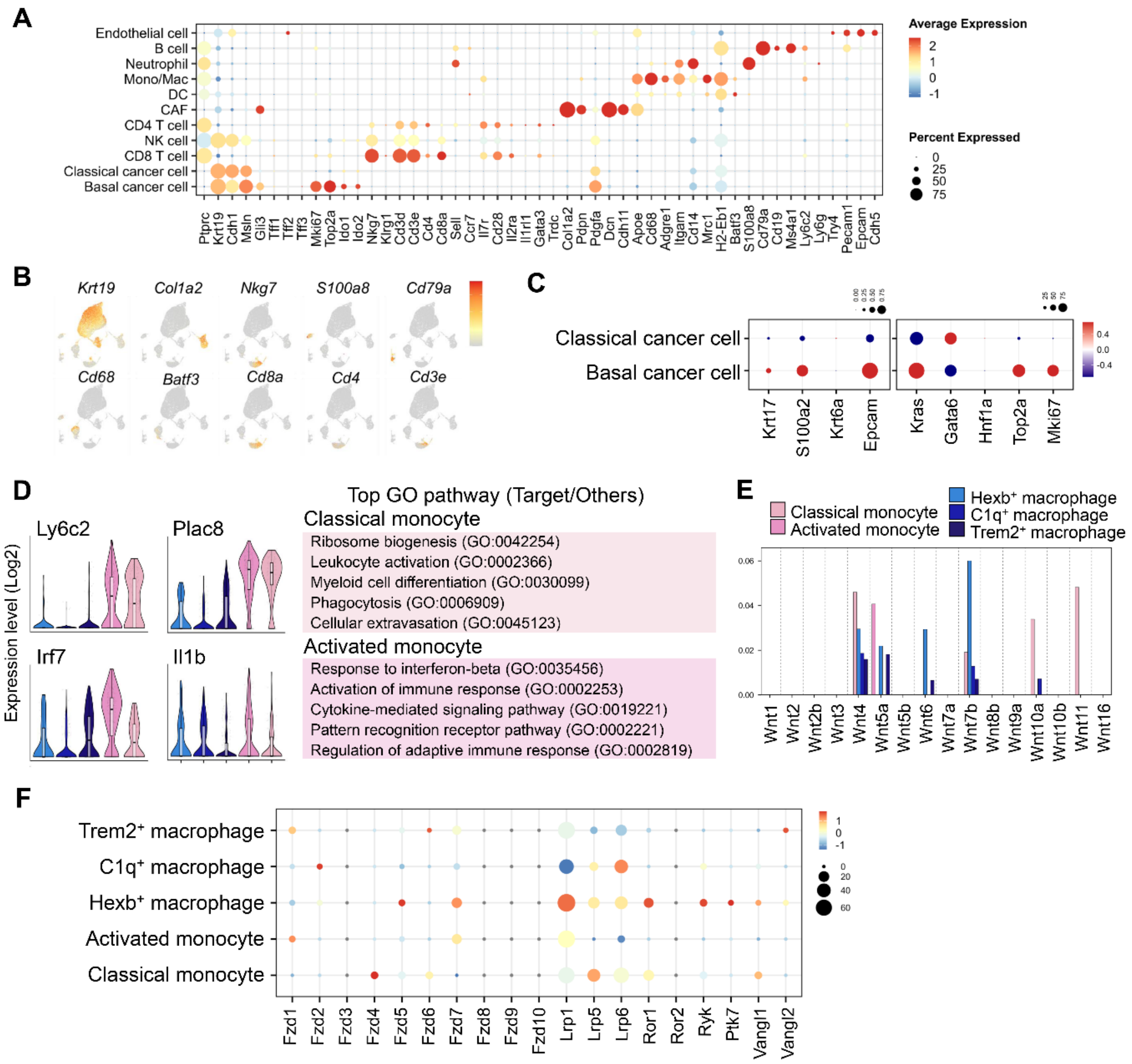

**Supplementary Figure S3. Single-cell annotation and WNT-related transcriptional features in the KPC tumor microenvironment.** (**A**) Dot plot showing expression of representative marker genes used to annotate major cell types in the mouse KPC tumor scRNA-seq dataset. Dot color indicates scaled average expression and dot size indicates the fraction of cells expressing each gene within the indicated cell type. (**B**) Feature plots showing expression of lineage markers used for cell type annotation. (**C**) Dot plots showing subtype marker expressions distinguishing classical versus basal-like cancer epithelial states. Left, epithelial identity/basal markers (e.g., Krt17, S100a2, Krt6a, Epcam); right, classical lineage and proliferation markers (e.g., Kras, Gata6, Hif1a, Top2a, Mki67). Dot color denotes scaled average expression, and dot size denotes percent expressed. (**D**) Violin plots depicting selected genes enriched in classical versus activated monocyte states (left), with top enriched GO terms for each monocyte state shown (right). (**E**) Bar plot summarizing Wnt ligand isoform transcript levels across mono/macrophage subsets. Bar indicates average gene expression. (**F**) Dot plot showing expression of WNT pathway receptors and transcription factor (TF) across monocyte/macrophage subsets, including Frizzled receptors (Fzd1–6), noncanonical receptors (Ror1/2), and canonical pathway components (Tcf/Lef family). Dot color indicates scaled average expression, and dot size indicates percent expressed.

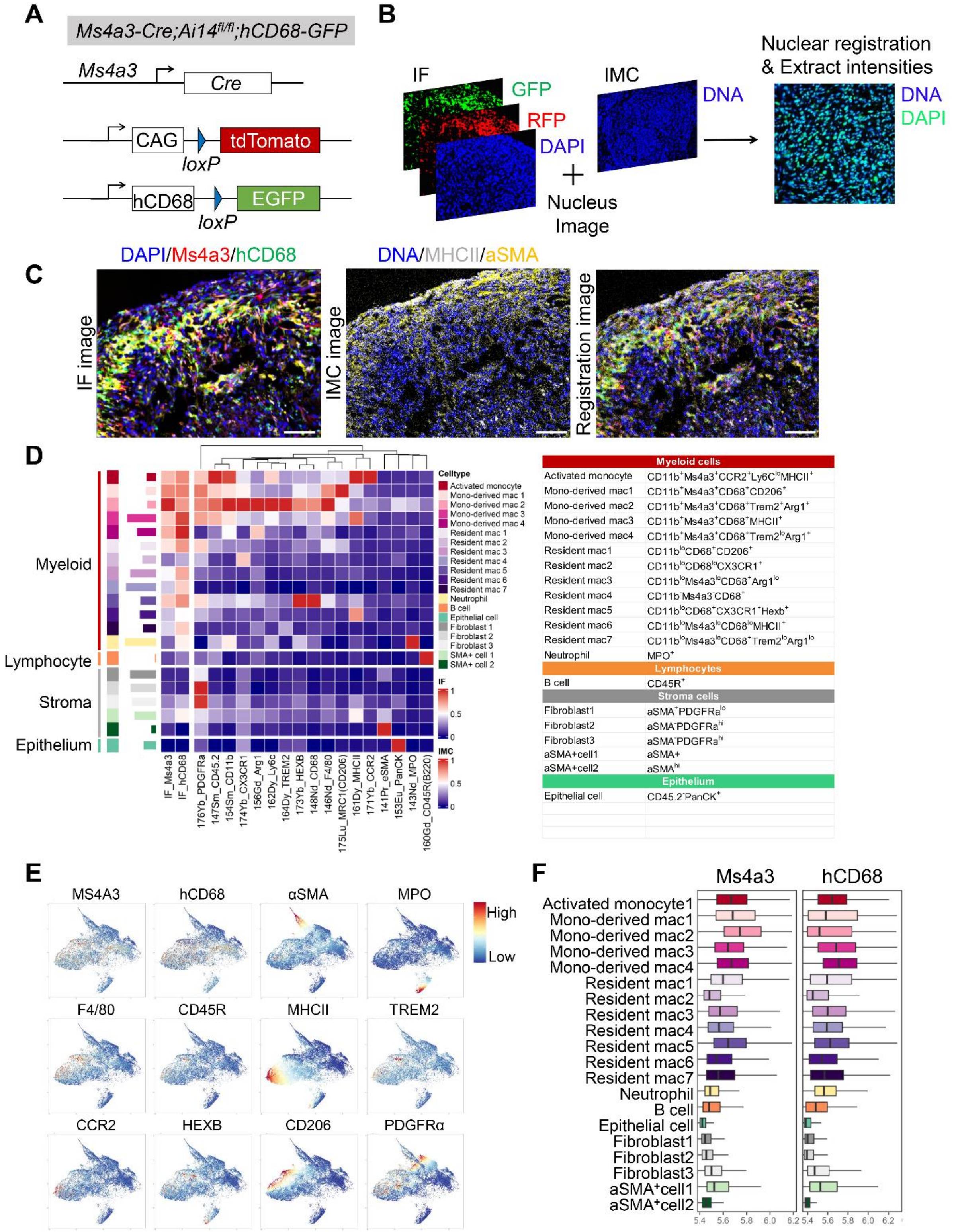

**Supplementary Figure S4. Validation of lineage-resolved IF–IMC integration and macrophage fate mapping.** (**A**) Schematic of the Ms4a3Cre-Ai14$^{\mathrm{Tdtomato}}$/hCD68-EGFP double-reporter mice strategy used to distinguish monocyte-lineage cells (Ms4a3 lineage tracing, tdTomato) and macrophages (hCD68-driven EGFP). (**B**) Overview of the image-registration pipeline used to align IF DAPI images with IMC DNA images, followed by nuclear registration and extraction of per-cell intensities for reporter channels. (**C)** Representative example of IF-IMC registration image, and the corresponding registered overlay demonstrating accurate spatial alignment between modalities. Scale bars, 100 μm. (**D**) Heatmap summarizing scaled marker intensities used for single-cell phenotyping across major compartments (left), with a detailed annotation schema shown at right for myeloid, lymphocyte, stromal, and epithelial populations (right). (**E**) UMAP feature plots of representative markers and reporter channels. (**F**) Box plots showing distribution of reporter intensities (Ms4a3 and hCD68) across annotated cell populations/subclusters, supporting classification of monocyte-derived versus resident-like macrophage states.

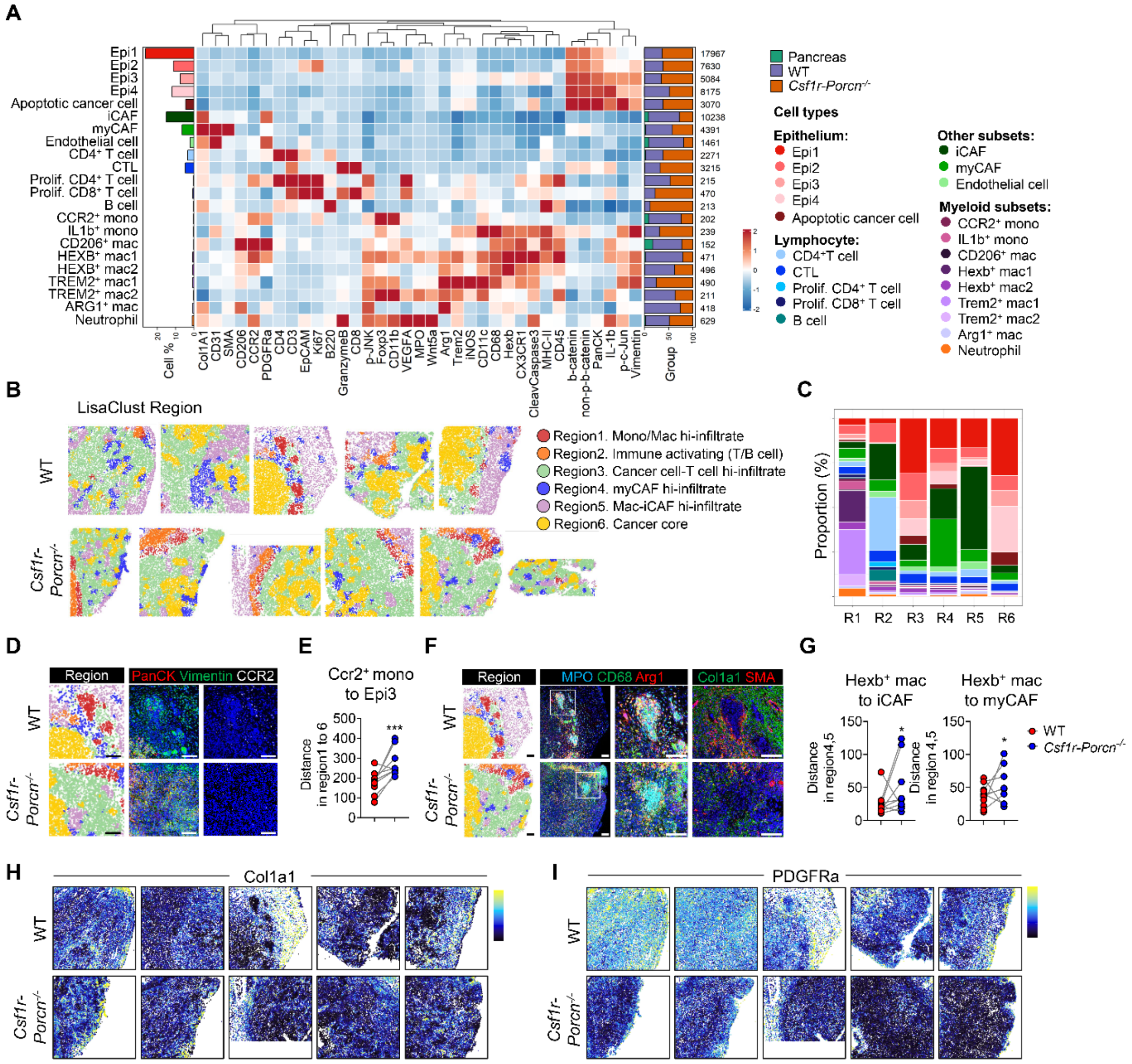

**Supplementary Figure S5. LisaClust-based spatial domain annotation and myeloid–stromal proximity analysis in PDAC.** (**A**) Heatmap summarizing scaled marker intensities used for single-cell phenotyping across major compartments. (**B**) Representative lisaClust region maps for WT and *Csf1r-Porcn*$^{-/-}$ tumors, with regions annotated as (1) monocyte/macrophage-high infiltrating niche, (2) immune-activating T/B-cell–rich niche, (3) cancer-core with T-cell high infiltration, (4) myCAF-high infiltrating niche, (5) macrophage–iCAF co-infiltrating niche, and (6) cancer-core niche. (**C**) Stacked bar plots quantifying the cell type proportion in each region. (**D**) Left, lisaClust region in WT and *Csf1r-Porcn*$^{-/-}$ Right, representative IMC images showing spatial relationships between tumor epithelium (PanCK), EMT cells (vimentin) and $CCR2^+$ monocytes. Scale bars, 100 µm. (**E**) Quantification of the mean distance between $CCR2^+$ monocytes and Epi3 tumor epithelial cells across regions 1. (**F**) Left, lisaClust region in WT and *Csf1r-Porcn*$^{-/-}$. Right, representative IMC images showing $MPO^+$ neutrophils, $CD68^+$ macrophages, ARG1 expression, and $Col1a1^+/\alpha SMA^+$ CAFs. Scale bars, 100 µm. (**G**) Nearest-neighbor distance between $Hexb^+$ macrophages and iCAFs (left) or myCAFs (right) within regions 4–5 in WT and *Csf1r-Porcn*$^{-/-}$ tumors. Marker expression overlay images for Col1a1 (**H**) and PDGFRα (**I**). Error bars indicate mean ± S.E.M.; Two-tailed unpaired t-test, * $p<0.05$, ** $p<0.01$, *** $p<0.001$, **** $p<0.0001$. n.s., not significant.

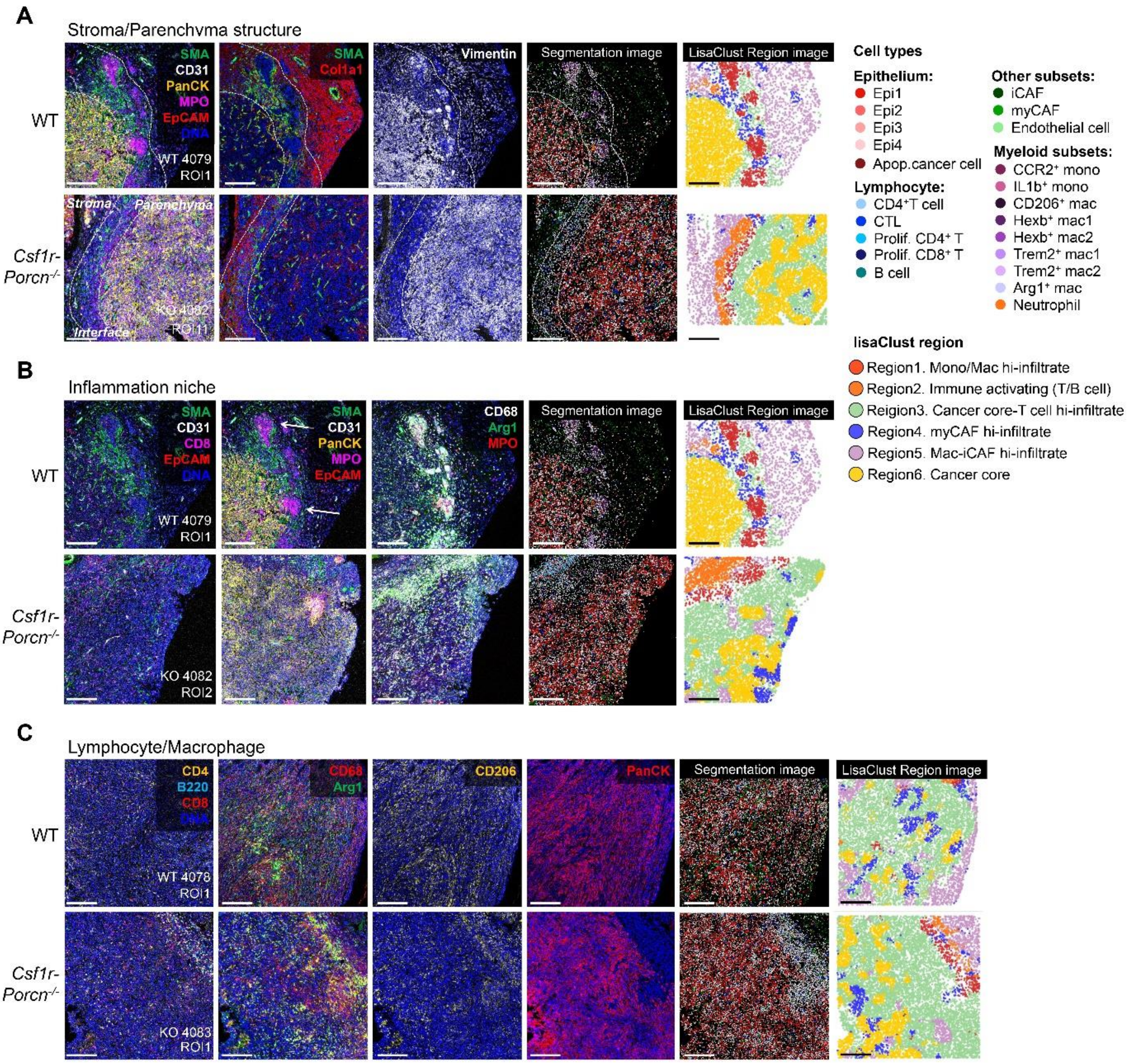

**Supplementary Figure S6. Representative IMC fields and LISA spatial maps in WT and *Csf1r-Porcn$^{-/-}$* KPC tumors.** (**A**) Four-color composite (left) highlighting stromal and epithelial compartments SMA (green), CD31 (white), PanCK (yellow), MPO (magenta), EpCAM (red), with an additional stromal composite SMA (green), Col1a1 (red) and Vimentin (white). Dashed lines indicate stromal–parenchymal boundaries. The same field is shown as a single-cell segmented image and as the lisaClust region map. (**B**) Marker composites emphasizing immune infiltration and myeloid localization, including SMA (green), CD31 (white), CD8 (magenta), PanCK (yellow), MPO (magenta), EpCAM (red) and a myeloidfocused composite CD68 (white), Arg1 (green), MPO (red). Single-cell segmentation and matched LisaClust region map are shown for the same ROI. (**C**) Immune-marker composite CD4 (yellow), B220 (cyan), CD8 (red), together with CD68 (red), Arg1 (green), CD206 (yellow), and PanCK (red) channels shown as indicated. Single-cell segmentation and the corresponding LisaClust region map are shown for the same field. Scale bars, 100 μm. ROIs shown are representative WT and *Csf1r-Porcn*$^{-/-}$ tumor regions; region color codes correspond to the LisaClust/LISA region definitions used in Fig. 5D-E
